# Population geometries in frontal cortex coordinate value-based decisions

**DOI:** 10.64898/2026.09.17.752402

**Authors:** Yonil Jung, Daigo Okada, Sarya Khandare, Rodrigo F.O. Pena, Hidehiko K. Inagaki

## Abstract

Decisions often involve trade-offs between benefits and costs, such as a small payment now vs. a larger payment after a delay. Although decision variables are distributed across frontal cortex, how these regions transform competing attributes into a choice remains unclear. Combining large-scale electrophysiology with causal perturbations as mice chose between different reward amounts and delays, we found that population geometry of decision variables differed across regions, reflecting distinct computations. Dorsal frontal areas, including motor cortex, collapsed competing attributes onto a decision axis shared across reward-delay contexts, with projection amplitude predicting choice probability; silencing these areas impaired action selection regardless of delay. In contrast, ventral prefrontal cortex (vPFC) represented choice in higher-dimensional subspaces organized by reward delay, whose geometry predicted how strongly delay influenced choice. Notably, inhibiting vPFC preserved the dorsal decision representation but selectively removed its modulation by reward delay, causing animals to behave as if delay no longer influenced choice. vPFC inhibition had no effect when mice chose between reward amounts. Frontal cortex therefore solves multi-attribute decisions through a generator–modulator architecture: dorsal frontal circuits maintain a shared choice axis guiding decisions across contexts, whereas vPFC maintains competing attributes in distinct subspaces that modulate decisions when options conflict.

## Introduction

Some decisions are straightforward: when two options differ only in payment, we simply choose the one that pays more. Decisions become more challenging when benefits and costs conflict, such as choosing between a small payment now and a larger payment later. How humans and animals behaviorally weigh competing benefits and costs—including reward, delay, effort, and risk—has been extensively characterized^1–3^, yet the neural mechanisms underlying these computations remain largely unknown.

To generate a choice, neural circuits must ultimately transform diverse task variables into action-selecting representations that guide behavior (Fig. 1a). These representations can be organized in different geometries within population state space, where each dimension corresponds to the activity of one neuron and each point represents the population state at a given moment^4,5^. Decision circuits may maintain contextual choice representations, in which choice axes differ across combinations of benefits and costs (Fig. 1b). Such a representation may function as a lookup table that flexibly maps each combination onto an action. Alternatively, they may maintain an abstract (generalizable) choice representation, in which a choice axis is shared across these combinations, allowing the same readout to guide choice across contexts (Fig. 1c). Such a representation could implement a common currency^6,7^ by compressing competing attributes into a scalar decision variable. Both types of representational geometry have been observed during decision-making, yet in largely separate studies^5,8–14^. Whether these distinct geometries reflect different computations across brain areas within the same animal, and how regions with different geometries causally contribute to behavior, remain unclear.

**Figure 1.**
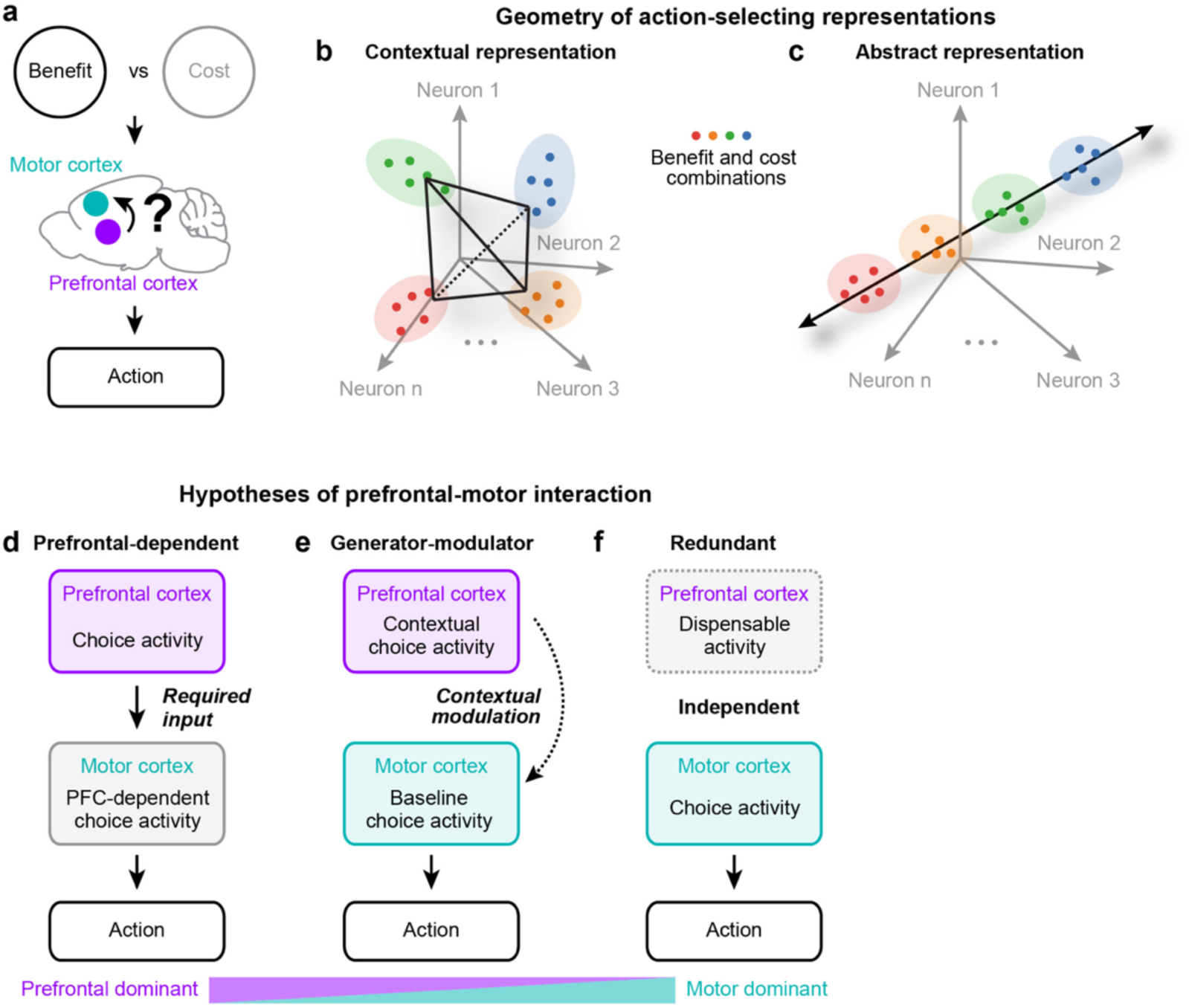
Representation geometry and frontal cortical interactions for value-based decision-making. **a,** Schematic of frontal cortical circuits integrating conflicting task variables (costs and benefits) to select an action. **b,** Contextual representational geometry mapping different combinations of task variables onto actions. **c,** Abstract representational geometry compressing task variables into a scalar value that guides action. Circle, population activity; Color, a distinct combination of benefit and cost. **d-f,** Possible configurations of interactions between PFC and MCx. Colored boxes denote the circuit component with an autonomous causal contribution; gray boxes denote dependent or dispensable activity; dotted outline denotes dispensable activity.

Answering these questions requires understanding how distinct frontal circuits contribute to decision computations. Frontal cortex has long been implicated in higher-order cognitive functions^8,15–18^, with prefrontal cortex (PFC) traditionally considered central to value-based decision-making^19–22^. Neurons throughout PFC encode reward, cost, value, and choice signals across species^6,23–30^, and perturbations of PFC impair value-based decisions^3,26,31^. Yet these computations must ultimately influence motor circuits to execute a choice. Notably, accumulating evidence indicates that motor and premotor cortex (MOp and MOs; MCx) are not merely passive output stages of PFC but also encode these decision variables and may contribute directly to decision-making^32–39^. This raises a central question: do prefrontal and motor cortices implement distinct computations reflected in different representational geometries, and if so, how do PFC computations shape motor cortical activity that ultimately drives choice?

We formalized a spectrum of circuit architectures that differ in how computations are distributed across frontal areas^15,40–43^. Under a *prefrontal-dependent* hypothesis (Fig. 1d), prefrontal areas provide signals required for motor areas to express an action-selecting representation. Prefrontal perturbation should therefore broadly disrupt choice across task contexts. Under a *generator–modulator* hypothesis (Fig. 1e), motor areas maintain an action-selecting representation independent of prefrontal input, whereas prefrontal areas selectively modulate motor decision activity according to task context, such as when benefits and costs conflict. Prefrontal perturbation should therefore selectively reduce context-dependent changes in choice while preserving choice generation itself. Under a *redundant* hypothesis (Fig. 1f), prefrontal and motor areas independently generate action-selecting representations, predicting little effect of prefrontal perturbation. Together, these hypotheses span a spectrum of motor-area autonomy, from complete dependence on prefrontal input to complete independence. These architectures can be associated with, but are not uniquely defined by, distinct action-selecting representational geometries, which we then measure directly.

To distinguish among these hypotheses, we developed a temporal discounting task in which head-restrained mice chose between two options whose reward amounts and delivery delays varied across blocks. The large number of trials (469 ± 77 trials per session; mean ± SD; 77 mice) enabled large-scale electrophysiology (24,765 cells, 52 mice) combined with causal perturbations, allowing us to characterize population geometry and quantify perturbation effects across frontal areas. Using these approaches, we identify a generator–modulator organization of frontal cortex: dorsal frontal areas (MCx and dorsal PFC) maintain a shared choice axis across contexts, whereas ventral prefrontal cortex (vPFC) maintains contextual representations that modulate dorsal choice activity to incorporate context-specific trade-offs, enabling multi-attribute decisions.

## Results

### Mice select actions based on both reward amount and delay

In the temporal discounting task, mice were presented with two lickports: one delivered a small, immediate reward (0–0.5 s from lick to water delivery), whereas the other delivered a large reward either immediately or after a delay of up to 4 s (Fig. 2a). Within each block of trials, the side associated with the large reward, reward magnitude, and reward delay were fixed, but these contingencies varied across blocks to assess their influence on choice (block duration randomly sampled from 5–25 trials; the large-reward side alternated across blocks; Fig. 2b; Extended Data Fig. 1-3). On each trial, an auditory go cue (3 kHz, 0.6 s) instructed animals to report their choice by licking the left or right spout. Because block transitions and reward contingencies were not cued, mice had to make choices based on recent choices and reward outcomes (i.e., trial history). Consequently, during the intertrial interval (ITI; 10.36 ± 2.41 s; mean ± SD), mice had to maintain information about recent trial history and/or prepare their upcoming choice before responding to the subsequent go cue.

**Figure 2.**
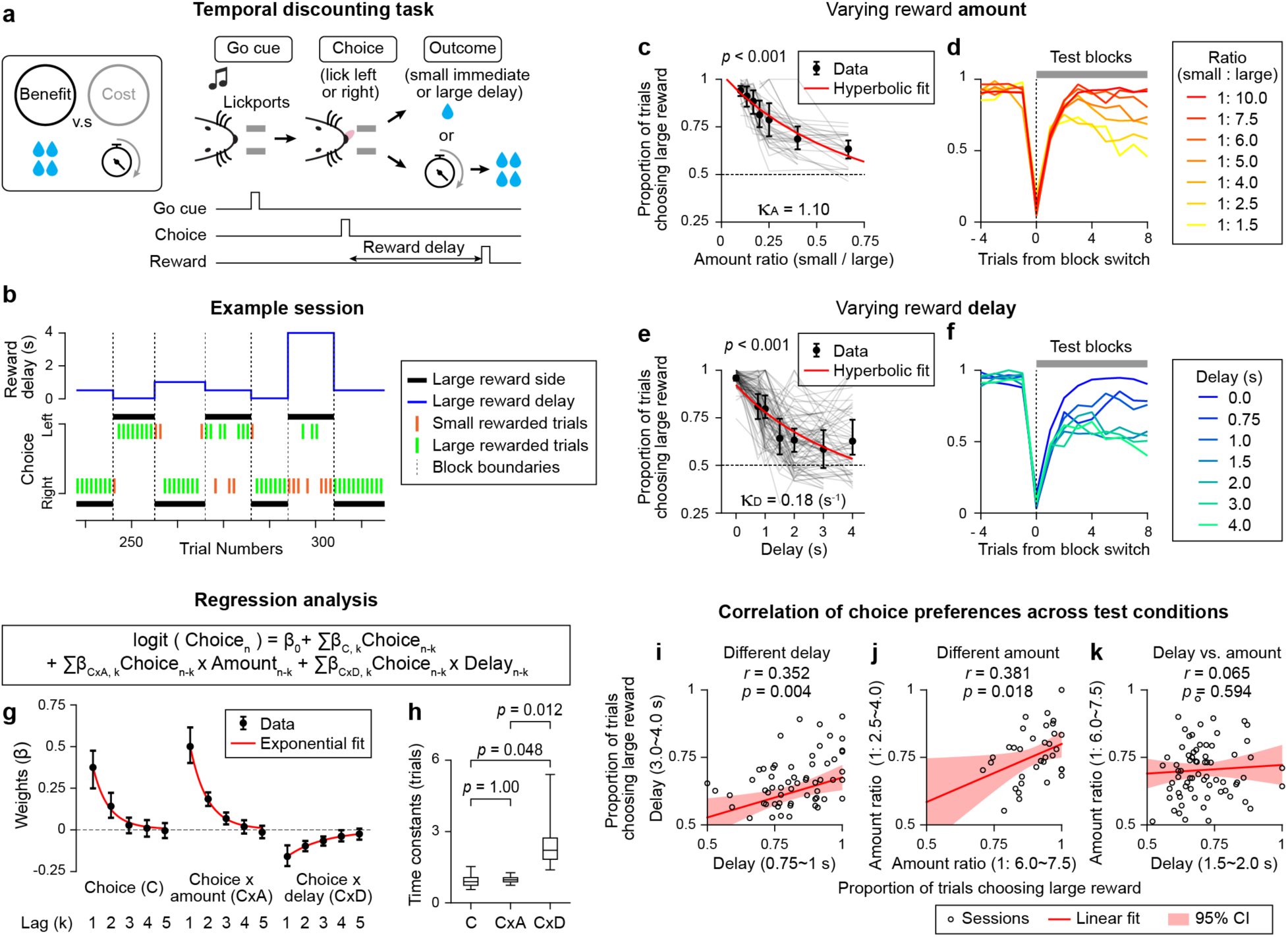
Behavioral performance in the temporal discounting task. **a,** Schematic of the temporal discounting task. **b,** Example behavioral session. Only a portion of the session is shown. **c,** Proportion of trials in which the large reward was chosen as a function of the reward amount ratio, with the large reward amount varied while delay was held constant. Line, individual session. Circle, mean; bar, 95% confidence interval (CI; hierarchical bootstrap); red line, hyperbolic fit; κ_A_, reward amount sensitivity coefficient (Methods); *p*-value, permutation test with the null hypothesis that the observed κ_A_ is equal to or greater than that obtained after trial-shuffling. n = 7 mice. **d,** Proportion of trials choosing the large reward around block switches. Line, mean. Color, amount ratio. **e-f**, same as in **c,d** for reward delay varied while reward amount was held constant. κ_D_, temporal discount coefficient (Methods). n = 7 mice. **g,** Regression coefficients of a logistic regression model explaining choice. Circle, mean; bar, 95% CI. red line, exponential fit; n = 6 mice. **h,** Time constants estimated from exponential fits to the regression weights in **g**. *: *p* < 0.05 (bootstrap followed by *Bonferroni’*s correction). In all boxplots across figures, central line, median; box edges, 25th and 75th percentiles; and whiskers the most extreme values within 1.5× the interquartile range. **i,** Correlation of choice preferences (proportion of trials choosing large-delayed reward) between shorter (0.75∼1 s) and longer (3∼4 s) reward delays. Circle, session. Red lines, linear regression. Shade, 95% CI. *r*, *Pearson’s* correlation coefficient. *p*-value, bootstrap testing the null hypothesis *r* is 0. n = 7 mice. **j,** Same as in **i,** comparing different large reward amount ratios. n = 7 mice. **k,** Same as in **i,** comparing varying reward delay vs. amount ratio. Reward amount and delay were varied within the same session to enable within-animal comparisons. n = 6 mice.

We first tested how reward amount shapes choice by randomly varying it across interleaved test blocks while holding reward delay constant (7 mice, 36 sessions; both small and large rewards delivered after a 0.5-s delay; Fig. 2c,d). Under a high reward amount ratio (small: large = 1:10), mice rapidly switched to the large-reward side following block transitions (1.35 ± 1.09 trials; mean ± SD) and maintained a strong preference for that side throughout the block (Fig. 2c,d). When the reward ratio was changed by reducing the large-reward amount, preference for the large-reward option decreased and reaction time increased (Fig. 2c,d and Extended Data Fig 1-2). Thus, mice flexibly adjusted their choices according to reward magnitude.

We next examined the effect of reward delay by varying the delay to large-reward delivery across blocks (0 – 4.0 s), while the small reward was delivered immediately (0-s) and the reward amount ratio was held constant (small: large = 1:4; 7 mice, 83 sessions; Fig. 2e,f). Increasing the delay reduced preference for the delayed large-reward option and increased reaction time (Fig. 2e,f and Extended Data Fig. 1-2). This delay-dependent decline in preference followed a quasi-hyperbolic profile, consistent with temporal discounting observed across species^2,44,45^. The estimated temporal discount rate, *κ*_D_ (0.18 ± 0.03 s^-1^; mean ± SEM; Methods), fell within the range previously reported for freely moving rodents^46–48^. Because longer delays also increase trial duration and reduce reward rate per unit time^49^, in some sessions we matched trial durations while varying delay. Mice still discounted delayed rewards, indicating that this effect could not be explained solely by differences in reward rate (Extended Data Fig. 3; n = 7 mice per condition).

The only information available to guide mice’s choices was recent trial history, including previous choice and outcomes (reward amount and reward delay). To quantify the trial-by-trial influence of these variables, we fit a logistic regression model to animals’ choices (Fig. 2g). The analysis revealed three major determinants of choice: (1) mice tended to repeat the same lick direction^50,51^ (2) larger rewards increased the probability of repeating the same choice on the next trial; and (3) longer reward delays reduced this probability (Fig. 2g and Extended Data Fig. 2h; see also Extended Data Fig. 4 for a fit to a Q-learning model^36,52–54^ yielding similar results).

Although both reward amount and reward delay influenced choice, two observations suggest that they contribute to decision through distinct computational processes.

First, regression analysis showed that reward delay influenced choice over 2.5 ± 1.1 trials (mean ± SEM; decay time constant; Methods), whereas the effects of previous choice and reward amount decayed substantially faster, with time constants of approximately one trial (Fig. 2h). Consistent with this notion, following a small, immediate reward trial, animals were more likely to switch choices in blocks where the large reward was delivered immediately (0-s delay block) than in blocks where the large reward was delayed (>1-s delay block; Extended Data Fig. 2g, left). Because the immediately preceding trial was identical in both conditions (small immediate reward), this difference indicates that animals retain information about reward delays over trials when guiding subsequent choices. In contrast, such multi-trial effects were not evident when reward amount ratio was manipulated (Extended Data Fig. 2g, right), consistent with reward amount exerting little influence beyond the immediately preceding trial.

Second, temporal discounting varied substantially across animals and sessions (Extended Data Fig. 2a,b). Despite this variability, sensitivity to reward delay was highly consistent: in sessions where mice discounted more strongly at one delay duration, they tended to discount more strongly at other durations (Fig. 2i). A similar consistency was observed for reward amount sensitivity (Fig. 2j). In contrast, delay and reward amount sensitivity were uncorrelated across animals (Fig. 2k), suggesting that they represent independent behavioral dimensions.

Together, these findings suggest that reward amount and delay are processed, at least in part, differently within decision circuits. We therefore examined how these variables are represented and geometrically organized across frontal cortex.

### Distributed coding but distinct temporal dynamics across frontal cortical areas

Using Neuropixels probes^55–57^, we recorded 12,426 neurons across frontal cortex and surrounding areas (71 sessions in 6 mice; Fig. 3a,b; the location of all recorded units registered to Allen common coordinate framework^58,59^; Extended Data Fig. 5). Across regions, individual neurons encoded diverse combinations of task variables. For example, many neurons exhibited persistent spiking activity during the ITI, anticipating upcoming choice. Some encoded the upcoming choice regardless of the reward delay on the preceding trial (Fig. 3c), whereas others exhibited choice-selective activity only following reward-delay trials (Fig. 3d).

**Figure 3.**
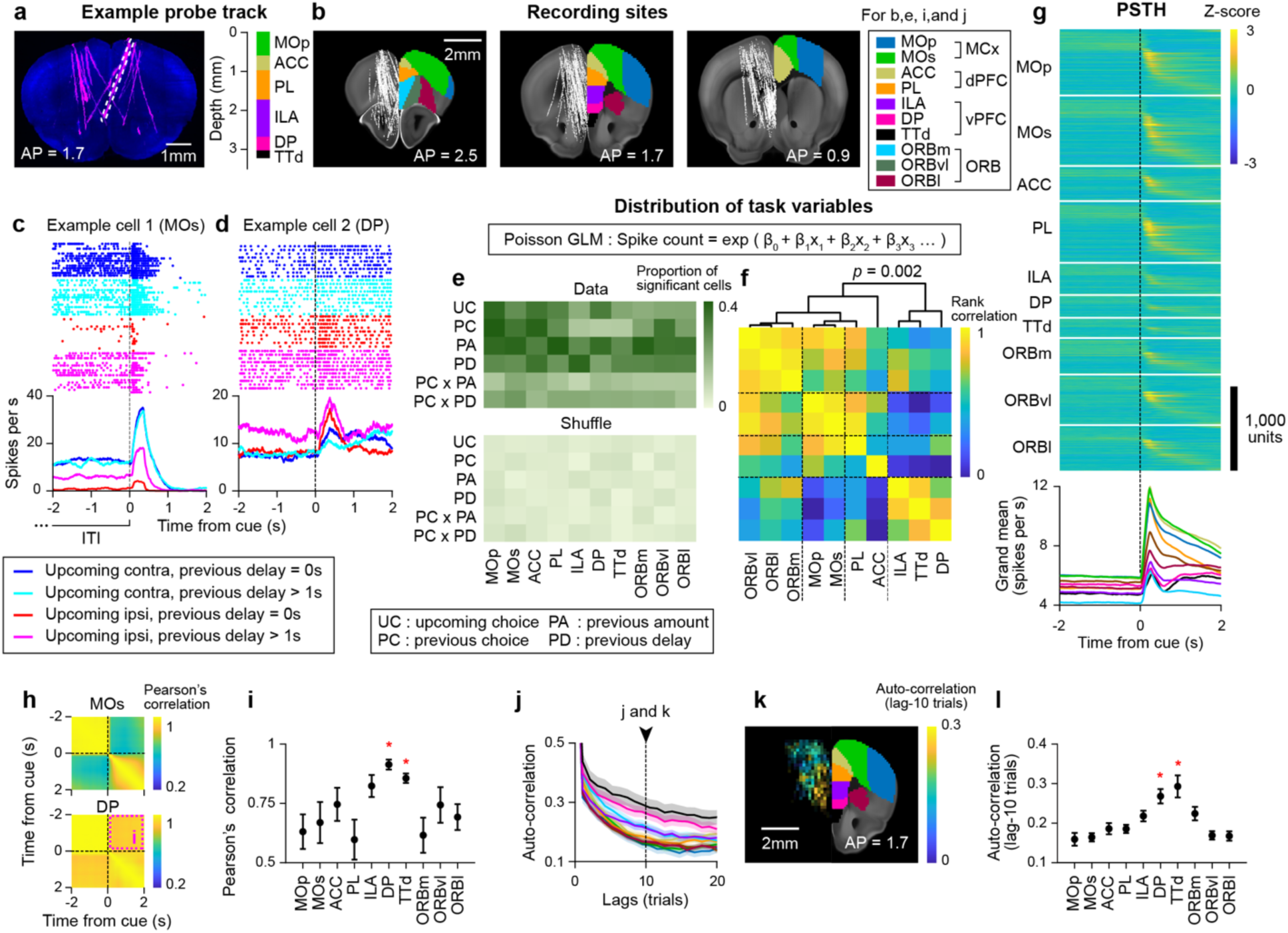
Spiking dynamics across frontal cortical areas. **a,** Left, light-sheet image of an example brain (coronal). Magenta, Neuropixels probe tracks; blue, autofluorescence. Right, corresponding brain regions traversed by the example probe track, indicated by white dashed lines. **b,** unit locations (white dots) registered to the Allen CCF (n = 16,136 units, 13 mice). **c-d,** Example cells. Top, spike raster. Trials are sorted by trial type (indicated by different colors). Bottom, peri-stimulus time histogram (PSTH). Contra/Ipsi, contralateral/ipsilateral relative to the recorded cell. **e,** Proportion of cells significantly encoding each task variable in each brain area (see **Extended Data** Fig. 6 for statistics). Top, data; bottom, trial shuffle control (Methods). **f,** Pairwise similarity of task-variable encoding profiles across brain areas, quantified by Spearman’s rank correlation across the proportions of significantly encoding cells shown in **e**. Dendrogram, hierarchical clustering based on encoding-profile similarity (Methods). *p*-value, bootstrap with a null hypothesis that vPFC is distinct from others (Methods). **g**, Top, Z-scored activity across frontal cortex. Neurons sorted by peak firing time in held-out trials. Bottom, grand average spike rate across areas. Color, same as in **b**. **h,** Pearson’s correlation matrix comparing MOs (DP) population activity across time points. Magenta dotted box, correlation quantified in **i**. **i,** Pearson’s correlation between activity before and after the cue. Circle, mean; Bar, SEM (bootstrap); *, *p* < 0.05, leave-one-region-out bootstrap test comparing each region with the mean of the remaining regions. **j,** Autocorrelation of ITI activity across brain areas. Line, mean; Shade, SEM (bootstrap). Color, same as in **b**. **k,** Spatial distribution of lag-10 autocorrelation**. l,** Comparison of lag-10 autocorrelation across brain areas. Circle, mean; Bar, SEM (bootstrap); * same as in **i**.

To quantify task representations across frontal cortex, we fit a Poisson generalized linear model to each neuron’s spike counts using six task variables: upcoming choice, previous choice, previous reward amount, previous reward delay, and two interaction terms between previous choice and either reward amount or reward delay. We applied recursive feature elimination with cross-validation to identify neurons that significantly encoded each task variable^60^.

During the ITI (−5 - 0 s before the go cue), neurons encoding each task variable were widely distributed across areas^61–63^ (Fig. 3e and Extended Data Fig. 6). Nearly all areas encoded all task variables significantly above shuffle control levels (α = 0.05 following Benjamini–Hochberg procedure), with the only exception being taenia tecta, dorsal part (TTd)^64^, which did not encode previous choice (Fig. 3e and Extended Data Fig. 6). Despite this broad encoding, hierarchical clustering of the encoding fractions revealed that the motor cortex (MCx; MOp and MOs), dorsal PFC (dPFC; PL and ACC) and orbitofrontal cortex (ORB; ORBm, ORBl and ORBvl) exhibited similar patterns of task-variable encoding, whereas the ventral PFC (vPFC; ILA, DP, and TTd)^64,65^ formed a distinct cluster (Fig. 3f). Thus, although nearly all regions significantly encoded all task variables, vPFC differed in the relative proportions of neurons representing each variable.

The temporal dynamics also differed in vPFC. First, population activity patterns in MCx remained stable throughout the ITI, as reflected by strong temporal correlations, followed by an abrupt transition after the go cue (Fig. 3h, top). This transition resembles reported shifts from motor planning to execution^66–68^. This cue-triggered transition in population activity pattern was accompanied by a sharp increase in spike rate following the go cue (Fig. 3g and Extended Data Fig. 6n) and was observed in MCx, dPFC, and ORB (with a weaker cue response in ORB; Extended Data Fig. 6m,n). In contrast, vPFC exhibited much weaker cue-triggered changes, reflected by higher temporal correlations and weaker cue-evoked increases in spike rate (Fig. 3h,i and Extended Data Fig. 6m,n).

Second, trial-to-trial autocorrelations of spiking activity during the ITI varied across regions. These time constants exhibited a topographic organization^69,70^, with neurons in vPFC generally exhibiting longer time constants than those in dorsal areas (Fig. 3j-l).

Together, these findings reveal a functional distinction between the vPFC and other frontal areas beyond task encoding patterns. Dorsal frontal regions (MCx and dPFC; ORB showed similar patterns but weaker cue responses) exhibited strong cue-triggered dynamics and short across-trial timescales, whereas vPFC exhibited weaker cue-triggered dynamics and longer across-trial timescales. These differences in task encoding and temporal dynamics may reflect different roles in value-based decision-making.

### Dorsal frontal activity maintains a graded decision representation and is required for choice

We hypothesized that dorsal frontal regions, with their trial-locked dynamics, support trial-by-trial action selection. To test this hypothesis, we asked whether 1) dorsal frontal activity is more predictive of upcoming choice than vPFC activity, and 2) perturbing dorsal frontal cortex impairs choice.

We first examined how strongly population activity in each frontal region predicted upcoming choice during the ITI. We trained a linear decoder (regularized logistic regression) to discriminate upcoming left versus right choices (Fig. 4a). To ensure that decoder performance reflected the upcoming choice rather than the previous choice, training trials were subsampled to balance previous choice (Methods). As reliable single-session decoding requires a sufficient number of simultaneously recorded neurons, we grouped frontal areas into four regions based on anatomical proximity, which broadly aligned with hierarchical clustering of their task-variable encoding profiles (Fig. 3f): MCx, dPFC, vPFC and ORB. Decoder performance was quantified on held-out trials using receiver operating characteristic (ROC) analysis, measured as the area under the curve (AUC).

**Figure 4.**
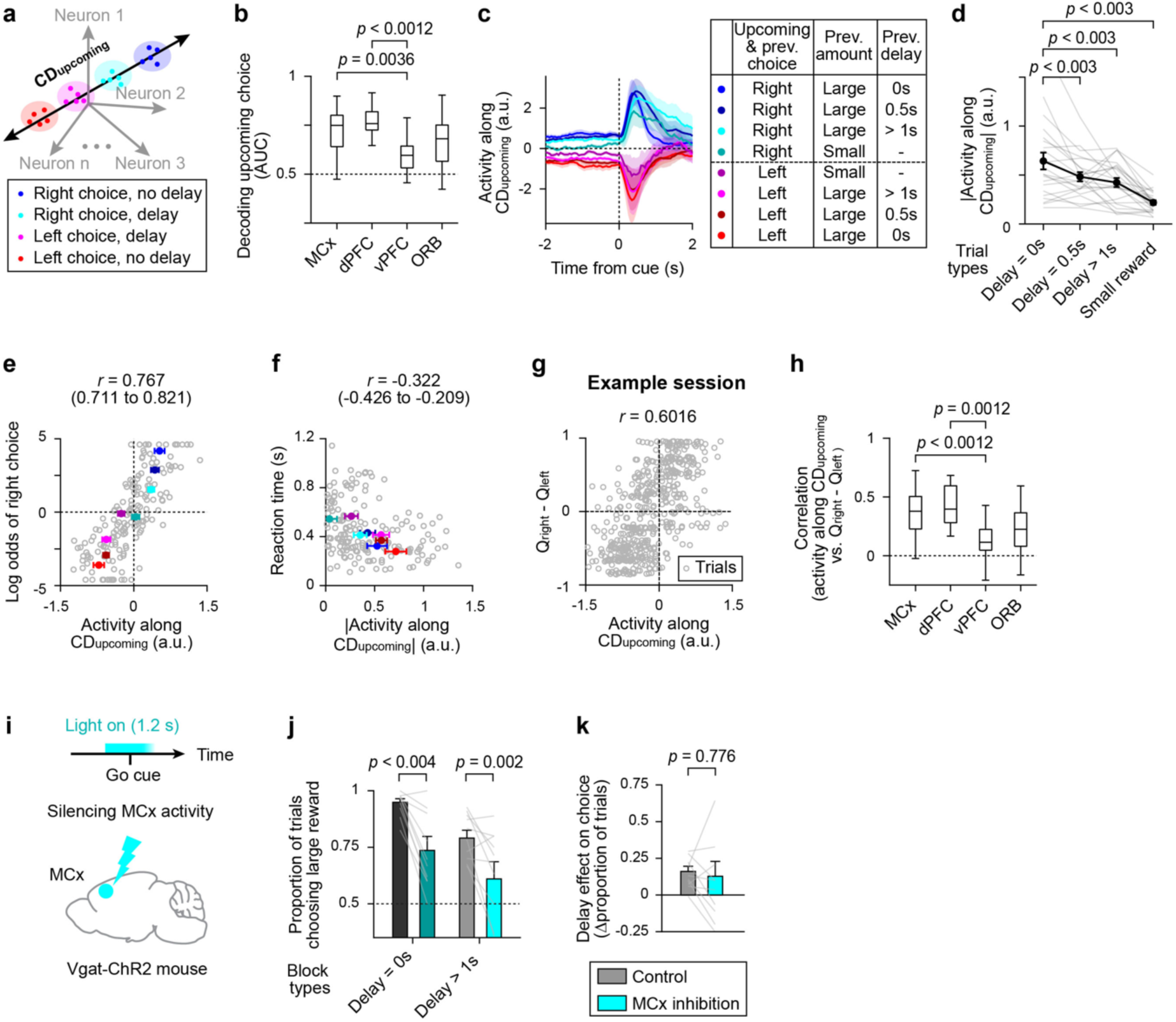
Graded decision coding across delay contexts in MCx. **a,** Schematic of **CD_upcoming_**. **b,** Session-wise decoding of upcoming choices in each brain area. Boxplots (**b** and **h**), same format as in Fig. 2g. *p*-value, hierarchical bootstrap comparing means with *Bonferroni* correction for six pairwise comparisons following a Kruskal-Wallis test (p = 5.60⨉10^-6^). All other pairs were *p*> 0.05. n = 31, 15, 27, 29 sessions for MCx, dPFC, vPFC, and ORB (**b-h**). See Extended Data Fig. 5h for the numbers of units, sessions, and mice. **c,** MCx population activity projected along **CD_upcoming_**. Color, trial type. Line, mean; shade, SEM. Sessions with a reliable decoding (AUC > 0.7) were included (same criterion for **c-f**). **d,** Absolute value of **CD_upcoming_** activity as a function of previous trial types. *P*-value, bootstrap test comparing the mean for Delay = 0 s vs. other conditions followed by *Bonferroni* correction for three comparisons. **e,** Relationship between MCx ITI **CD_upcoming_** activity (stay trials only) and the log odds of rightward choice across trial-history types (including both stay and switch trials). For projection, only stay trials were included to avoid confounding of activity by choice, as graded mean activity could arise solely from differences in behavioral choice probability. Gray circle, session; Colored circle, mean; Bar, SEM. **f,** Same format as in **e** for reaction time. **g,** Relationship between MCx ITI **CD_upcoming_** activity and the estimated difference in Q values between the two choices in an example session. Circle, trial. **h,** Session-wise Pearson’s correlation between **CD_upcoming_** activity and QR-QL in each brain area (as in **g**). *p*-value, hierarchical bootstrap comparing means with *Bonferroni* correction for six pairwise comparisons following a Kruskal-Wallis test (p = 3.02⨉10^-6^). All other pairs were *p* > 0.05. **i,** Schematic of bilateral MOs silencing in Vgat-ChR2 mouse. **j,** Proportion of trials choosing the large reward with MOs silencing. Bars, mean; error bars, SEM (bootstrap); gray line, individual mouse (n = 10 mice). *p* values, hierarchical bootstrap comparing means with *Bonferroni* correction for two comparisons. **k,** Change in the delay effect (the difference in the proportion of trials choosing the large reward between Delay = 0 s and Delay > 1 s) with MOs silencing. Same format as in **j**.

Upcoming choice was decoded significantly better from MCx and dPFC than from vPFC, whereas ORB showed intermediate decoding performance (Fig. 4b; see Extended Data Fig. 5h for the number of analyzed cells, sessions, and mice per area). Mean spike rate and Fano factor were similar across areas, ruling out differences in basic firing properties as an explanation for the observed decoding differences (Extended Data Fig. 7a-e). Moreover, the superior decoding performance of MCx relative to vPFC remained significant after accounting for the number of recorded neurons (Extended Data Fig. 7f). Together, these results show that population activity predicts upcoming choice more accurately in dorsal frontal areas (MCx and dPFC) than in vPFC.

We next examined the structure of activity along the decoder axis during the ITI, which we refer to as **CDupcoming**. **CDupcoming** activity in MCx remained separated according to upcoming choice throughout the ITI (Fig. 4c; blue vs. red traces indicate right-vs. left-lick trials). Although **CDupcoming** was defined to maximally discriminate upcoming left versus right choices, activity along this direction was not simply categorical. Notably, even when animals selected the same action on consecutive trials (stay trials), the amplitude of **CDupcoming** activity varied systematically with the previous reward outcome (Fig. 4c,d and Extended Data Fig. 7i-k): it was largest following a large reward delivered without delay, decreased with increasing reward delay, and was lowest following a small reward. Across trial-history conditions (defined by previous choice, reward amount, and delay), variation in **CDupcoming** activity (calculated only from stay trials to control for differences in choice composition) paralleled differences in choice probability and reaction time calculated across all trials (Fig. 4e,f)^34^. Together, these findings show that activity along the population direction that discriminated upcoming choices represented choice in a graded manner that scaled with choice probability, rather than categorically, during the ITI.

Under decision policies such as the softmax rule, choice probability is a monotonic function of the underlying action-value difference^53^. As expected from its relationship with choice probability, **CDupcoming** activity correlated on a trial-by-trial basis with the estimated action-value difference (Qright−Qleft), inferred by fitting a Q-learning model to behavioral data (Fig. 4g,h). Together, these results are consistent with activity along **CDupcoming** representing a graded decision variable underlying choice.

In contrast, a coding direction derived from post-cue activity (0–0.5 s after the go cue) also discriminated left versus right choices but exhibited a bimodal representation that was largely independent of previous trial type (Extended Data Fig. 7o-q), possibly reflecting its closer relationship to motor output. Thus, MCx activity transitioned from a graded representation of upcoming choice probability during the ITI to a categorical representation of the selected action after the go cue.

We next tested whether dorsal frontal activity, particularly in MCx, is causally required for choice. To this end, we used transgenic mice expressing ChR2 in GABAergic neurons (Vgat-ChR2-EYFP mice^71^) with clear-skull preparations^72^ (Methods). We bilaterally silenced MCx centered on the orofacial region, anterior lateral motor cortex (ALM^72–74;^ anterior 2.5 mm lateral 1.5 mm from Bregma), during a 1.2-s window around the go cue by scanning a blue laser across the cortical surface (beginning 0.6 s before the go cue onset; 488 nm, 1.5 mW per spot; <20% of randomly interleaved trials; 10 mice; Fig. 4i; Methods). Simultaneous recordings confirmed reliable suppression of MCx spiking activity without significantly affecting vPFC spiking activity (Extended Data Fig. 8a-h; as in ^72,75^).

Silencing MCx impaired choice, reducing preference for the large reward without significantly affecting the no-lick rate or reaction time (Fig. 4j and Extended Data Fig. 8). This implies that MCx is required for maintaining and/or selecting a choice in this behavior, consistent with its activity patterns (Fig. 4b-h) and the established role of MCx in motor planning^72,76,77^. Importantly, the behavioral effect was comparable in no-delay (0-s) and >1-s delay blocks, indicating that MCx activity is required for choice regardless of delay context (Fig. 4j,k). By contrast, the distinct temporal dynamics and task-variable encoding of vPFC raise the possibility that it contributes to decision-making through a different computation.

### Distinct population geometries between dorsal vs. ventral frontal areas

The lower decodability of upcoming choice in vPFC may reflect, at least in part, differences in representational geometry rather than weaker choice information. If choice-coding directions differ across behavioral contexts, such as different block types, a single decoder trained across pooled trials will be poorly aligned with the coding direction in each context and therefore perform poorly^5,78–80^.

To test this possibility, we performed single-session decoding analyses conditioned on delay block type (with 0-, 0.5-, or >1-s delay). For each session, we trained a linear decoder to discriminate choice using ITI activity from trials within a given block type and then tested its performance on held-out trials from either the same (within-condition) or a different (cross-condition) block type. Because switch and small-rewarded trials were rare in the 0- and 0.5-s delay blocks, we trained decoders using only stay trials following large rewards. This ensured that differences across block types could be attributed to reward delay rather than differences in trial composition. Decoder performance was compared with circular-permutation controls that preserved trial-to-trial autocorrelation (Methods). A decoder trained on one block type should generalize to other blocks if choice is encoded along a shared direction across delays (‘abstract coding’; Fig. 5a, top). In contrast, cross-condition decoding should fail if the choice-coding direction differs across block types (‘contextual coding’; Fig. 5a, bottom).

**Figure 5.**
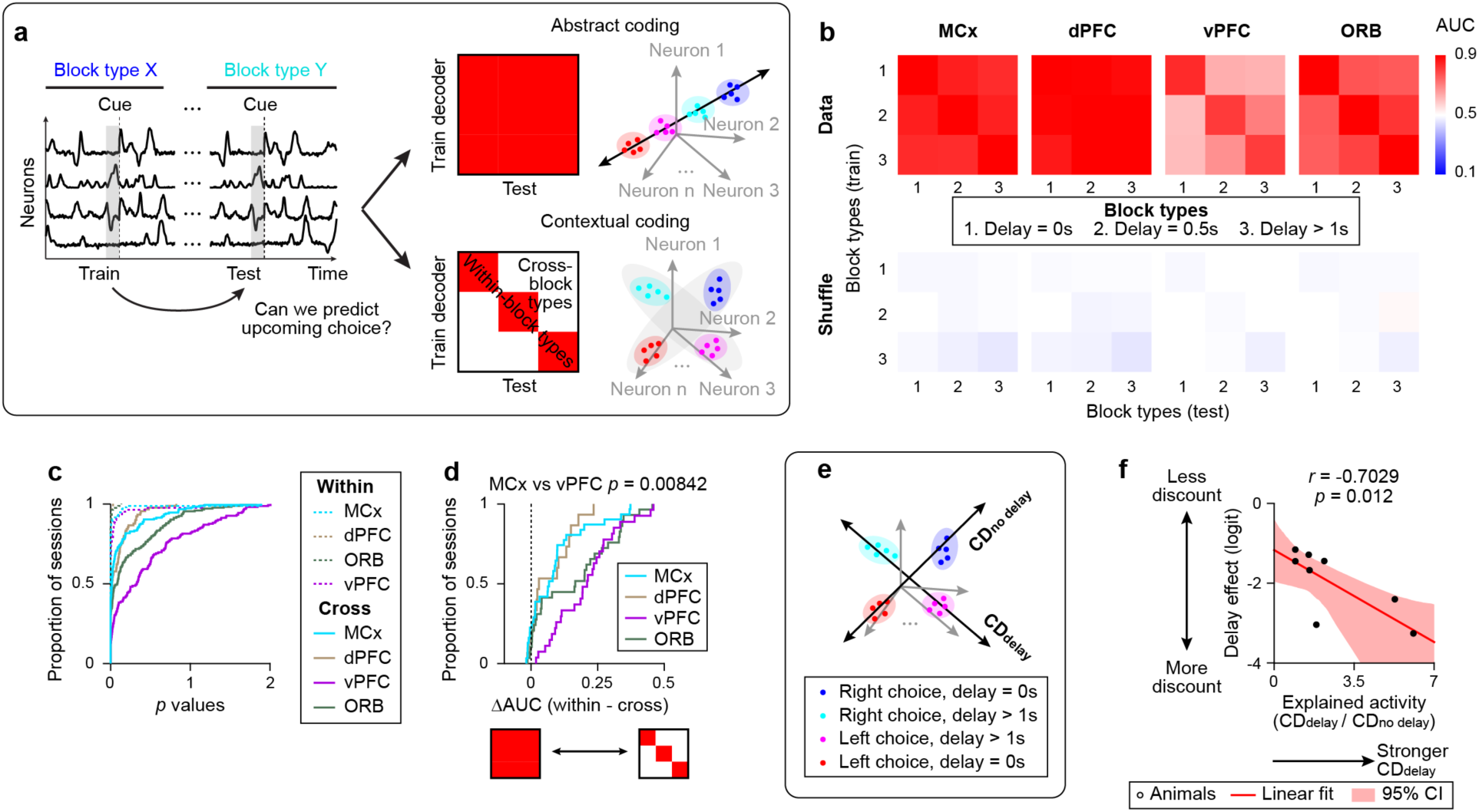
Representation geometry in frontal cortex. **a,** Schematic of the decoding analysis. A decoder trained on one block type was evaluated on held-out trials from the same or the other block type. High generalization (top) indicates abstract coding, whereas low generalization across block types (bottom) indicates high-dimensional contextual coding. **b,** Decoding analysis across brain regions. Top, mean AUC across sessions; bottom, mean AUC in circular permutation controls. n = 31, 15, 27, 29 sessions for MCx, dPFC, vPFC, and ORB (**b-d**). **c,** Distribution of *p* values across sessions (permutation test comparing AUC in data vs. permutated data in **b**; Methods). Dashed line, within-block-type comparison. Solid line, cross-block-type comparison. **d,** Cumulative distribution of the difference between within-block and cross-block decodability across sessions. *P*-value, Kolmogorov–Smirnov test followed by *Bonferroni* correction for six comparisons. **e,** Schematic of **CDdelay** and **CDno delay**. **f,** Ratio of squared activity explained by **CDdelay** and **CDno delay** versus the behavioral delay effect (logit). Circle, animal. *r*, *Pearson*’s correlation; *p*-value, a bootstrap test for a nonzero correlation.

Decoders generalized robustly across block types in MCx and dPFC (Fig. 5b), consistent with abstract coding and the shared choice axis (Fig. 4). In contrast, vPFC showed little cross-condition generalization, whereas ORB exhibited an intermediate level of generalization (Fig. 5b). These results are consistent with contextual coding and with the reduced performance of vPFC in the single **CDupcoming** analysis (Fig. 4).

All regions, including vPFC, exhibited reliable within-condition decoding (Fig. 5c, dotted lines). However, cross-condition decoding performance declined (Fig. 5c, solid lines), with the reduction being significantly greater in vPFC than in MCx (Fig. 5d). Several controls indicate that this regional difference cannot be explained by sampling or temporal confounds. First, the greater reduction in cross-condition decoding in vPFC persisted when analyses were restricted to sessions with high within-condition decoding accuracy (AUC > 0.9), ruling out poorly defined decoders as an explanation (Extended Data Fig. 9a,b). Second, the same dissociation was observed in sessions with simultaneous MCx and vPFC recordings, demonstrating that the difference reflects regional properties within the same animals (Extended Data Fig. 9c,d). Third, it was not explained by differences in the number of recorded cells (Extended Data Fig. 9e).

Finally, decoders generalized better to temporally distant trials within the same block type than to temporally closer trials from a different block type (Extended Data Fig. 9f). Thus, the reduced cross-condition decoding in vPFC cannot be explained by the temporal proximity of within-condition training and test trials.

We observed a similar regional difference in pseudopopulation analyses, which enabled finer area-specific comparisons by pooling neurons across sessions, albeit at the cost of single-trial information. These analyses revealed contextual coding (distinct choice-coding directions in the 0-vs. >1-s delay blocks) in vPFC (ILA/DP/TTd), ORBm, Striatum, and NAc (Extended Data Fig. 9h,i). Notably, Striatum and NAc are subcortical regions that receive projections from vPFC^65,81^. These findings indicate that, in contrast to the abstract representation of choice observed in MCx and dPFC, choice information in vPFC and its downstream circuits is represented in a context-dependent manner.

Notably, whether coding is abstract or contextual depends on the represented variable. First, dorsal frontal areas (MCx and dPFC) were not globally lower-dimensional^82^, as their population dimensionality, quantified by participation ratio in neuronal space, was comparable to that of vPFC and ORB (Extended Data Fig. 9l). Moreover, decoders of previous reward delay (or amount) trained on one choice (e.g., whether the left choice had a delay vs. no delay) failed to generalize to the other choice, even in MCx and dPFC (Extended Data Fig. 9m,n), indicating that abstract coding is selective for choice (right vs. left) across delay contexts rather than a general property of population geometry. Importantly, contextual coding in vPFC was not simply a consequence of manipulating a reward attribute that altered choice. Although varying the reward amount ratio influenced choice similarly to varying reward delay (Fig. 2c,d), choice coding was abstract across reward ratios in vPFC, as reflected by the high cosine similarity of choice-coding directions (Extended Data Fig. 9j,k).

Altogether, during the ITI, dorsal frontal regions (MCx/dPFC) represent choice along a shared direction, whereas vPFC represents the same information in a context-dependent geometry spanning distinct choice-coding directions. Importantly, abstract versus contextual coding was specific to the represented variable and behavioral context, rather than a general property of population geometry. These findings raise the possibility that population geometry is shaped by the computational demands of the task: a shared choice-coding direction in MCx/dPFC may facilitate cross-condition action selection, whereas contextual representations in vPFC may support delay-context-dependent differences in choice bias.

If contextual geometry in vPFC supports computations involving reward delay, then variation in this geometry should predict delay-context-dependent differences in choice behavior. Contextual coding in vPFC involves distinct coding directions for each block type: **CDno delay** and **CDdelay** for 0- or >1-s delay, respectively. We quantified the relative prominence of these coding directions by calculating the proportion of population activity captured by **CDdelay** and **CDno delay** in their corresponding contexts (Methods). The ratio of these values predicted differences in temporal discounting across animals: greater relative activity along **CDdelay** was associated with greater temporal discounting (Fig. 5f; n = 8 mice). This relationship suggests that vPFC geometry reflects behavioral sensitivity to reward delay.

### vPFC contextual coding shapes delay-dependent choice and MCx activity

Having established that vPFC choice-coding geometry is context-dependent and predicts temporal discounting, we next asked whether these contextual representations are causally required for delay-dependent choice and how they interact with the choice representation in MCx.

Correlation analysis revealed context-dependent coupling between vPFC and MCx. Specifically, activity along **CDdelay** in vPFC was most strongly correlated with the MCx **CDupcoming** activity in delay blocks (>1-s delay), whereas activity along **CDno delay** was most strongly correlated with MCx **CDupcoming** activity in no-delay blocks (0-s delay; Extended Data Fig. 10). Thus, vPFC activity projected onto each context-specific coding direction was more strongly coupled with MCx activity in the corresponding context. This dynamic coupling is consistent with the ‘Prefrontal-dependent’ and ‘Generator–Modulator’ architectures, although common inputs could also contribute to the observed correlations in the ‘Redundant’ architecture (Fig. 1d-f).

To distinguish between these architectures, we next optogenetically inhibited vPFC while simultaneously recording from MCx (Fig. 6a). Under the ‘Prefrontal-dependent’ architecture, silencing vPFC should eliminate or markedly reduce MCx **CDupcoming** activity, impairing choice across block types (Fig. 6i, left). Under the ‘Generator–Modulator’ architecture, silencing vPFC should selectively reduce the delay-context-dependent modulation of MCx **CDupcoming** activity and choice (Fig. 4d,e), while preserving the baseline **CDupcoming** activity and choice (Fig. 6i, middle). Under the ‘Redundant’ architecture, vPFC silencing should have little effect on either MCx **CDupcoming** activity or behavior (Fig. 6i, right).

**Figure 6.**
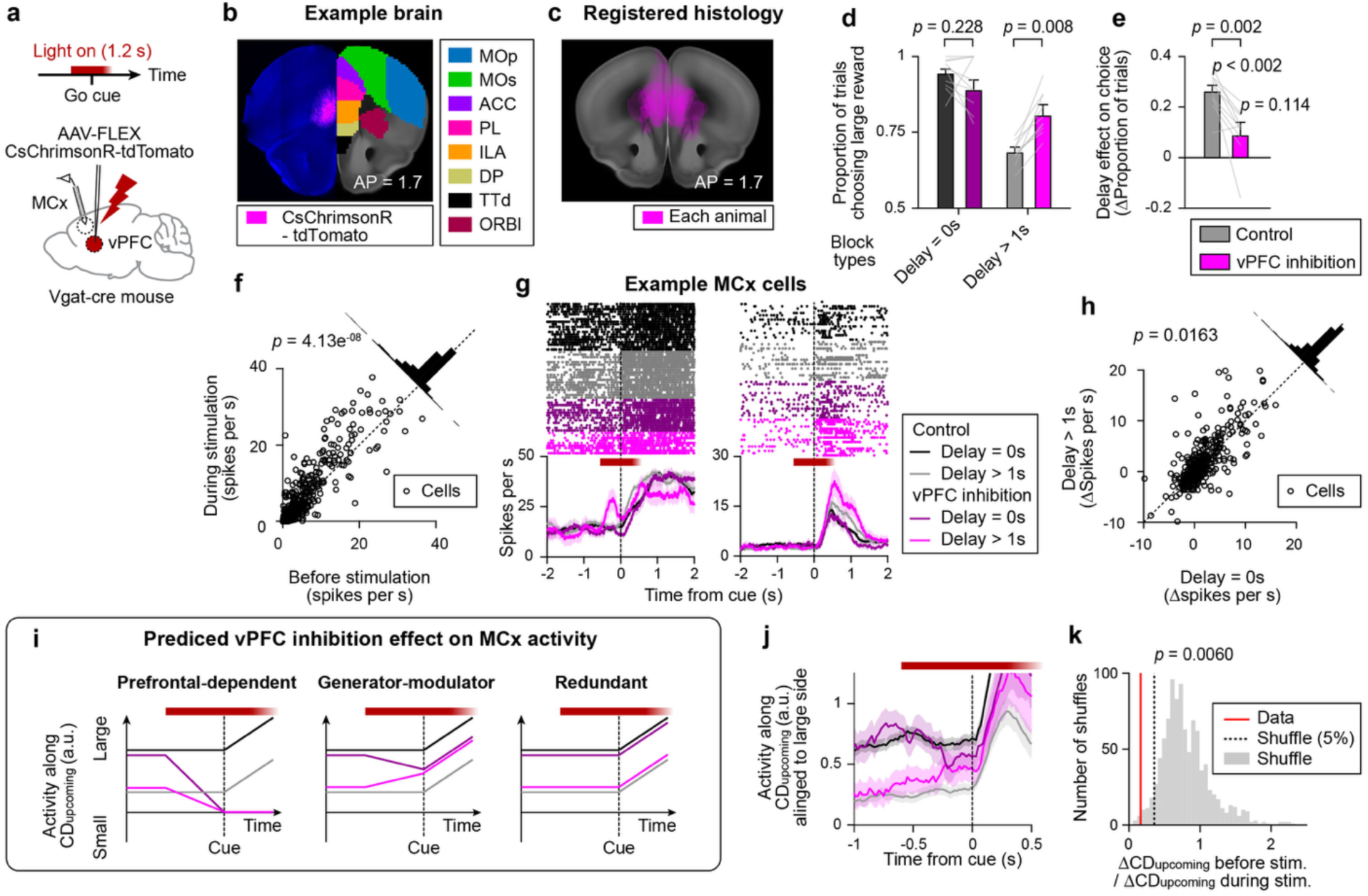
Transient optogenetic perturbation of vPFC. **a,** Bilateral vPFC inhibition in vGAT-ires-Cre mouse expressing ChrimsonR-tdTomato. **b,** Example brain. Blue, autofluorescence; Magenta, ChrimsonR-tdTomato. Right, Allen CCF. **c,** Virus expression overlay across mice (n = 10 mice). **d,** Proportion of trials choosing the large reward with vPFC inhibition (n = 10 mice). Same format as in Fig. 4j. **e,** Change in the delay effect with vPFC inhibition. Same format as in Fig. 4k. **f,** Spike rate of MCx putative pyramidal neurons before and during vPFC inhibition. Top right, histogram of spike rates during minus before stimulation. n = 564 neurons; *p-*value, signed-rank test. **g,** Example MCx cells. Top, spike raster. Trials are sorted by trial type (indicated by different colors). Bottom, PSTH. **h,** Change in spike rate during vPFC inhibition in delay vs. no-delay blocks. The same format as in **f**. Stay trials following large rewards were analyzed. **i,** Schematics of predicted vPFC inhibition effects under different hypotheses (Fig. 1d**-f**). **j,** MCx population activity projected along **CD_upcoming_**. The sign of **CD_upcoming_** activity was standardized so that positive values corresponded to the large-reward side, allowing pooling across blocks with different large-reward sides. Line, mean; shade, SEM. **k,** Permutation test examining whether vPFC inhibition reduces the difference in **CD_upcoming_** activity between delay and no-delay blocks. Bars, distribution of permutations (1000 iterations); red line, data. **g-l,** n = 437 neurons, 6 mice.

We achieved bilateral transcranial photoinhibition of vPFC by virally expressing the red-shifted opsin CsChrimsonR^83^ (AAV2/5-hSyn-FLEX-CsChrimsonR-tdTomato) in vGAT-ires-Cre mice, thereby activating local GABAergic neurons with red light (637 nm, 5 mW per spot; Fig. 6a-c; Methods). This approach avoided implant-related damage to dorsal frontal areas while preserving access for simultaneous silicon probe recordings. Recordings from vPFC during photostimulation confirmed increased spiking in putative GABAergic neurons and suppression of putative pyramidal neurons, validating this approach (Extended Data Fig. 11).

We bilaterally inhibited vPFC for 1.2 s around the go cue (beginning 0.6 s before cue onset) in randomly interleaved trials (<30%) while mice performed the task (n = 10 mice). Inhibition significantly increased preference for the large-reward option during delay blocks (>1-s delay) but showed a trend toward decreased preference during no-delay blocks (0-s delay), effectively eliminating the reward-delay-dependent difference in choice, which was no longer significantly different from zero (Fig. 6d,e). In contrast, although varying reward amount altered choice behavior to a similar extent as varying reward delay in control trials, vPFC inhibition had no effect when mice chose between reward amounts (n = 8 mice; Extended Data Fig. 12b). Moreover, perturbing ORB during reward-delay manipulations produced no detectable effect on choice (n = 6 mice; Extended Data Fig. 12c). ORB lies anterior to vPFC and exhibits less delay contextual coding (Fig. 5b-d), supporting the anatomical specificity of the vPFC effect. Together, these results indicate that vPFC is not generally required for value-based decision-making but is specifically required for integrating reward-delay context into choice.

Next, we examined how optogenetic inhibition of vPFC altered MCx activity during behavior. vPFC inhibition significantly increased spiking activity in 173/564 of MCx neurons and decreased activity in 112/564 (signed-rank test, α = 0.05; n = 10 mice). On average, MCx firing rates increased by 1.58 ± 0.22 spikes per s (mean ± SEM; Fig. 6f, g), indicating that vPFC exerts a net suppressive influence on MCx activity. Notably, MCx neurons exhibited significantly larger increases in firing rate during vPFC inhibition in >1-s-delay blocks than in 0-s-delay blocks (0.49 ± 0.13 spikes per s; mean ± SEM; Fig. 6h), indicating that functional interactions between vPFC and MCx are stronger during delay than no-delay conditions. This difference was not observed during reward-amount manipulation or ORB manipulation (n = 8 mice; Extended Data Fig. 12l and p), indicating that the effect was specific to reward delay and vPFC.

We examined the mean spike rates in vPFC and MCx on control trials and found no significant differences between the 0- and >1-s delay blocks (Extended Data Fig. 7c). This argues against a simple model in which vPFC is more active during >1-s delay blocks and more strongly suppresses mean MCx activity. Instead, the delay-dependent difference in the strength of vPFC–MCx interactions may be specific to coding directions.

We therefore examined how vPFC inhibition altered MCx activity along **CDupcoming**, defined using control trials. If vPFC provides delay-context information to MCx, disrupting vPFC activity should reduce the context-dependent modulation of **CDupcoming** activity. Because photoinhibition was bilateral and our goal was to isolate the effect of reward delay, we pooled blocks in which the left or right spout was associated with the larger reward. Before photoinhibition onset, the separation of **CDupcoming** between delay and no-delay blocks was comparable in control and inhibition trials. Following vPFC inhibition, however, this separation was significantly reduced (Fig. 6j,k and Extended Data Fig. 13d-f). Thus, vPFC selectively supports the context-dependent modulation of **CDupcoming**, explaining its selective behavioral effect on delay-dependent choice.

Although vPFC inhibition selectively reduced the context-dependent modulation of MCx activity, MCx maintained robust **CDupcoming** activity that continued to predict upcoming choice during the inhibition (Extended Data Fig. 13c). Thus, vPFC is not required to maintain MCx choice-coding representation itself during the late ITI, arguing against the ‘Prefrontal-dependent’ architecture during this period (but these results do not exclude a role for vPFC in establishing this representation earlier in the ITI). Instead, vPFC continues to selectively provide delay-context information to an otherwise intact decision representation in MCx, modulating choice according to the current delay context. Together, these findings support the ‘Generator–Modulator’ architecture.

## Discussion

Value-based decision-making requires integrating benefits and costs into a single action. Such computations may be implemented through distinct neural algorithms associated with different population geometries, ranging from context-specific, lookup-table-like representations to a shared choice axis that implements a common currency for action selection. Although value-related signals have been reported throughout the frontal cortex, several fundamental questions remained unresolved: how representations are organized across regions, how regions with distinct population geometries interact, and how they causally contribute to decisions.

Our results reveal that frontal regions differed less in the information they encoded than in how this information was geometrically structured and utilized at the population level. Although reward, delay, and choice signals were broadly encoded across the frontal cortex, their population-level organization differed markedly across regions. Dorsal frontal regions, particularly MCx, serve as an essential substrate for choice formation and/or maintenance, as silencing this region impaired choice across delay conditions. At the population level, MCx maintains a shared choice axis during the ITI that predicts subsequent choice probability across delay contexts. This MCx activity may function as a common currency, integrating previous actions and outcomes into a graded decision signal from which the likelihood of each choice is computed through a softmax-like transformation^53,84^. Following the go cue, this graded action-value-like representation transitions into a categorical representation of the selected action, consistent with the transformation from decision formation to action execution.

In contrast, vPFC contains context-dependent representations during the ITI that selectively encode choice according to the delay context. Simultaneous recordings from MCx during vPFC inhibition revealed that vPFC is required for context-dependent modulation of both MCx population activity and choice behavior.

Together, these findings indicate that frontal cortex partitions computation across areas with distinct population geometries. Dorsal frontal circuits (MCx and dPFC) house shared choice-coding direction that supports choice across contexts, whereas vPFC circuits provide contextual choice representations that flexibly modulate those representations. This division was selective for reward delay: although manipulations of both reward amount and delay altered choice, only delay produced contextual geometry in vPFC, and only delay-dependent choice was sensitive to vPFC perturbation. Thus, the generator–modulator organization may be particularly important when competing reward attributes favor different actions, requiring contextual information to adapt decision-making.

Notably, this proposed division of labor closely parallels the established roles of prelimbic (PL; part of the dPFC) and infralimbic (ILA; part of the vPFC) cortex in fear conditioning. PL is required for the expression of conditioned fear across contexts, whereas ILA is critical for the context-dependent modulation of fear responses, such as during extinction learning^85,86^. DP/TTd have also recently been implicated in fear extinction, similar to ILA^64^. Although anatomical homologies between rodent and primate prefrontal cortex remain debated^87,88^, similar modulatory roles for prefrontal cortex have been proposed in primates^43,89,90^. Together, these parallels raise the possibility that the generator–modulator organization identified here reflects a general computational principle of frontal cortex rather than a mechanism specific to value-based decision-making in rodents.

Just as spatial navigation requires information about an animal’s location in physical space, flexible decision-making may require information about its location in “context space” defined by recent experience. Our results suggest that vPFC provides such contextual representations, allowing the same action to be prepared differently depending on behavioral context. The context-dependent coding observed in vPFC may arise from its stronger interactions with hippocampal circuits relative to dorsal regions^65,91,92^, or from distinct learning rules that promote the formation of context-specific representations. Notably, striatum and NAc, major projection targets of vPFC, exhibited similar contextual coding, raising the possibility that vPFC influences MCx through basal ganglia (NAc)–thalamocortical circuits^93,94^ or via its direct projection^65^.

One way to view this distinction is that vPFC performs a form of representational expansion, separating decision variables according to behavioral context, whereas MCx extracts a common action variable. Similar representation expansion motifs, which can facilitate the separation of overlapping inputs, are used in machine learning, and have been observed in other brain areas such as the cerebellar granule cell layer and the hippocampus^95–98^. Together, these findings raise the possibility that the combination of contextual and abstract representations constitutes a general computational strategy for flexible behavior across brain systems.

Decision-making varies substantially across individuals, particularly in situations involving conflict, where there is often no objectively correct answer and preferences are inherently subjective^93^. Our results show that the context-dependent population geometry in vPFC predicts differences in delay discounting across animals. This observation is consistent with studies showing that population activity can account for individual differences in choice and reaction times^99–101^. These findings suggest that behavioral variability may arise, at least in part, from differences in neural geometry—that is, in how decision variables are organized within high-dimensional population activity. Such geometries may be shaped through learning and accumulated behavioral experience^102–104^, providing a neural substrate that enables similar tasks to be performed using distinct behavioral strategies across individuals.

Together, our findings highlight the importance of population-level analyses, as such geometric features emerge from the collective organization of neural activity and are not readily apparent at the level of individual neurons. These results provide a unifying framework linking population geometry, causal circuit function, and behavioral flexibility: frontal cortex resolves value conflict through a generator–modulator architecture that combines contextual representations in vPFC with abstract decision representations in MCx.

## Acknowledgments

We thank Lorenzo Fontolan, Ryoma Hattori, Yue Liu, and Inagaki lab members for comments on the manuscript, M. Inagaki, M. Watabe, and L. Walendy for animal training, K. Shirley and A. Ilchenko for imaging, and H. Shearin and other MPFI ARC members for animal care.

## Author contributions

Conceptualization: HKI, YJ; Investigation: YJ, DO, SK, RP, HKI; Funding acquisition: HKI; Supervision: HKI; Writing – original draft: YJ, HKI; Writing – review & editing: YJ, HKI

## METHOD DETAILS

## EXPERIMENTAL MODEL AND SUBJECT DETAILS

### Mice

This study is based on both adult male and female mice (age > P60). We used three mouse lines: C57Bl/6J (JAX #000664), Vgat-ires-cre (JAX #28862)^105^, and VGAT-ChR2-EYFP (JAX #14548)^71^.

All procedures were in accordance with protocols approved by the MPFI IACUC committee. We followed the published water restriction protocol^106^. Mice were housed in a 12:12 reverse light: dark cycle and behaviorally tested during the dark phase. A typical behavioral session lasted between 1 and 2 hours. Mice obtained all of their water in the behavior apparatus (approximately 0.6 ml per day). Mice were implanted with a titanium headpost for head fixation and single-housed. Craniotomies for recording were made after behavioral training.

### Viral Injection

To virally express CsChrimsonR in vPFC, we followed published protocols (dx.doi.org/10.17504/protocols.io.bctxiwpn) for virus injection. AAV2/5 Syn-FLEX-CsChrimsonR-tdTomato^83^ (titer: 7.8×10^12; UNC Gene Therapy Vector Core) was injected into AP 1.7 mm, ML ± 0.5 mm, DV 2.7 mm, 100 nl in Vgat-ires-cre mice. To target ORB, coordinates of AP 2.5 mm, ML ± 0.5 mm, DV 2.5 mm were used.

### Behavior

At the beginning of each trial, an auditory cue consisting of three pure tones (3 kHz, 150 ms duration, 100 ms inter-tone interval, 74 dB) signaled the onset of a 3 s answer period, during which mice chose by licking either the left or right lickport. Lick was measured by detecting the contact of the tongue with the lickport using an electrical lick detector. One lickport delivered a small reward after a short delay (0 or 0.5 s from lick to water delivery), whereas the other delivered a large reward either immediately or after a delay of up to 4 s. The lickport associated with the large reward alternated in a block schedule, with block lengths randomly selected between 5 and 25 trials. There was no external cue indicating the block switch or reward amount/delay. If mice did not lick during the 3 s answer period, the trial would end without a reward (“no lick trials”). Trials were separated by an inter-trial interval (ITI), defined as the period from reward delivery to the onset of the next go cue. ITI consisted of a fixed offset and two random intervals: an interval sampled from an exponential distribution (*τ* = 1-s; Maximum, 4 s) to prevent animals from predicting cue onset, and a stop-licking period sampled from an exponential distribution ( *τ* = 2-s, Maximum 7 s) during which any lick restarted the stop-licking period to encourage lick suppression during the ITI. Together, these components resulted in a mean ITI of 10.36 s (minimum, 5.22 s; maximum, 21.03 s; Extended Data Fig. 3d). In approximately 3% of randomly interleaved trials, the auditory cue was omitted to assess spontaneous lick rate (‘no cue’ trials), and these trials were excluded from analyses. No water reward was delivered in no cue trials.

For training, mice were initially trained with 0.5-s reward delay (for both the small- and large-reward options) across all blocks to learn to alternate licking between the left and right lickports following the large reward. The reward delay was then introduced in every third block and progressively increased up to 1.5 s.

During behavioral testing, every third block served as a test block, in which the reward amount, reward delay, or both were randomly selected from predefined ranges. These test blocks were used for the behavioral analyses shown in Fig. 2 and Extended Data Fig. 1-3. For the varying-delay test, each test block was followed by a 0.5-s delay block (0.5-s delay for both small- and large-reward options) and then a no-delay block (0-s delay for both reward options), allowing paired comparisons between delay and no-delay blocks. For the varying-amount-ratio test, reward delay was fixed at 0.5 s for both small- and large-reward options across all blocks.

In Fig. 4, we recorded spiking activity while varying reward amount and reward delay across test blocks, and activity in all block types was included in the analyses. For single-session analyses in Figs. 4-6, we alternated among 0-, 0.5-, and >1-s (either 1.25-, 1.5- or 2-s) blocks to increase the number of trials available for analysis at each delay duration. To avoid human bias, the behavior was automatically controlled by Bpod (Sanworks) and custom MATLAB codes.

### Optogenetics

Photostimulation was deployed on < 30% of randomly selected trials in no-delay or delay blocks. To prevent mice from distinguishing photostimulation trials from control trials using visual cues, a ‘masking flash’ (1 ms pulses at 10 Hz) was delivered using 470 or 627 nm LEDs (Luxeon Star) throughout the trial. For ChR2 and CsChrimsonR, we used a 488 nm and 637 nm laser (OBIS 488 - 150C and OBIS LX 637 nm, Coherent), respectively. To avoid rebound excitation, we ramped down the laser power linearly during the last 300 ms of 1.2-s photostimulation^107,108^. As the behavioral effect weakened across successive sessions (consistent with adaptation observed in many chronic perturbation studies^109,110^), we analyzed only the first session for both behavior and electrophysiology.

### Extracellular electrophysiology

A small craniotomy (diameter, 0.5 - 1 mm) was made over the recording sites one day before the first recording session. Extracellular spikes were recorded using 64-channel two-shank silicon probes (Cambridge Neurotech), Neuropixels probe 1.0, or Neuropixels probe 2.0 (imec). Probes were acutely inserted in each session. For 64-channel probes, voltage signals were multiplexed, recorded on a PCI6133 board (National Instruments), and digitized at 400 kHz (14-bit). The signals were demultiplexed into 64 voltage traces sampled at 25 kHz and stored for offline analysis. All recordings were made with the open-source software SpikeGLX (http://billkarsh.github.io/SpikeGLX/). During recordings, the craniotomy was immersed in a cortex buffer (125 mM NaCl, 5 mM KCl, 10 mM glucose, 10 mM HEPES, 2 mM MgSO4, 2 mM CaCl2; adjust pH to 7.4). Brain tissue was allowed to settle for at least five minutes before recordings. Probe tracks labeled with CM-DiI were used to determine recording locations.

### Histology

Mice were perfused transcardially with PBS, followed by 4% PFA / 0.1 M PBS. Brains were post-fixed overnight. To reconstruct recording tracks, we cleared the brain followed by light-sheet microscopy. To clear the brain, we used the EZ Clear method^111^.

## QUANTIFICATION AND STATISTICAL ANALYSIS

### Behavioral analysis

We analysed the lick direction (choice) and time of the first lick (reaction time) after the cue onset in each trial. The no-lick rate was calculated as the probability of mice not responding within 3 s after the cue. To analyze behavior while mice were actively engaged in the task, we included data only after mice reached a large-reward choice rate of at least 65% across five 0.5-s-delay blocks and until they either failed to respond for five consecutive trials or developed a sustained side bias, defined as choosing only one lickport across four consecutive blocks. Blocks were excluded from behavioral and neural activity analyses if the lickport position was adjusted or if water was provided outside the task structure to re-engage an animal that had stopped responding, which rarely occurred during experiments. Trials without a response or cue presentation were excluded from analyses.

In the absence of a cue signaling a block switch, animals could not infer the switch until they experienced the first outcome following it. Therefore, the first trial after a block switch was considered part of the preceding block for analysis. Similarly, animals could not infer the test block type until they experienced the first large-reward outcome; therefore, we analyzed only trials following that outcome (except for experiments in which the small reward amount was altered: Extended Data Fig. 1l).

#### Fit to hyperbolic function and estimation of κ (Fig. 2c,e and Extended Data Fig. 1-3)

The proportion of trials in which animals selected the large-reward option as a function of reward delay or reward-amount ratio was fit with a hyperbolic function.

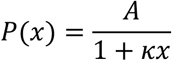

where A is a scaling parameter and *κ* quantifies sensitivity to reward delay (or reward ratio). Parameters were constrained to be non-negative and estimated by minimizing the residual norm between observed and predicted choice probabilities using a bounded Nelder–Mead simplex optimization. Confidence intervals for *κ* were estimated using a hierarchical bootstrap. Statistical significance (Fig. 2c,e. *p*-value) was assessed with a permutation test with the null hypothesis that the observed values were equal to or smaller than those obtained after trial shuffling.

#### Trial-history regression analysis of choice (Fig. 2g,h and Extended Data Fig. 2h)

To quantify how previous choices and their outcomes influenced subsequent decisions, we fitted a logistic regression model predicting the upcoming lick direction from the choice, reward amount, and reward delay experienced on preceding trials. Analyses included 71 behavioral sessions from 6 mice (31,806 trials). Trials without a response or cue presentation were excluded. Within each session, reward amount and reward delay were z-scored.

For each trial, the design matrix contained three groups of regressors: previous choice, previous choice × reward amount, and previous choice × reward delay, each evaluated over the preceding 10 trials (30 regressors in total). The probability of a rightward lick was modeled using logistic regression with L2-regularization. The regularization strength was determined by five-fold cross-validation on the pooled dataset and fixed for all subsequent analyses. To estimate the variability of the regression coefficients, we performed hierarchical bootstrapping by resampling animals with replacement and then resampling sessions with replacement within each animal (1,000 iterations). In each iteration, the model was trained on one half of the trials and evaluated on the held-out half. Trial splits were stratified according to the type of the preceding trial to preserve their relative proportions.

To quantify how the influence of previous trials decayed with time (Fig. 2h), regression weights from the first five lags were fitted with a single exponential function,

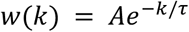

where *A* is the amplitude, *k* is trial, and *τ* is the decay time constant. Fits were performed independently for each regressor family and each resampling iteration using nonlinear least-squares optimization.

#### *Fit with a Q learning model* (Fig. 4g,h and Extended Data Fig. 4)

Choice behavior was described with a variant of Q-learning^53^ that incorporates both exponential forgetting of past action values and hyperbolic discounting of delayed rewards. The Q-learning framework describes how an agent learns to make decisions over time. Separate state–action values were maintained for the right-(Qright) and left-port (Qleft) options, which were updated according to whether each option was chosen or unchosen:

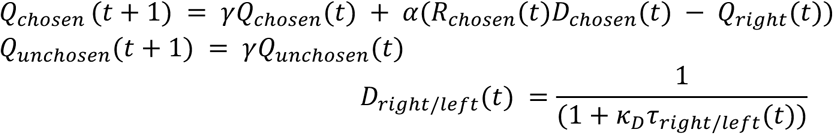

where *γ* is the forgetting factor, controlling how much past information is retained (0 for no forgetting, 1 for no memory decay), *α* is the learning rate, *R* is the actually delivered reward, and *D* is the discount factor, calculated from hyperbolic fit with discounting rate (*κ_D_*) and actual delay duration (*τ*).

Action selection followed a soft-max rule that also included a perseveration parameter (*ρ*) that depends on the previous lick direction (*a*(*t* − 1)):

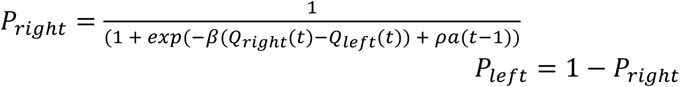

All Q-values were initialized to zero for the first two trials. Parameters were obtained separately for each session by maximizing the log-likelihood of the observed sequence of choices under the model with random restarts. For each fitted session we calculated the trial-by-trial choice probability and plotted it together with the 5-trial-smoothed lick direction trace. Predictive accuracy was quantified as the coefficient of determination (R²) obtained when regressing the smoothed empirical choice sequence onto the model-derived choice probability. To aid interpretation, Q-values, delays and reward magnitudes were linearly rescaled into non-overlapping y-axis bands.

### Histology analysis

We followed the previously described protocol to map the recording tracks to the Allen Common Coordinate Framework (CCF)^59,112^. The area with virus expression was quantified from binarized fluorescence images of coronal brain sections (Fig. 6c and Extended Data Fig. 12a-c). In Extended Data Fig. 12d, the AP location was defined by the section with the largest expression area. The center of viral expression was estimated as the center of the best-fitting ellipse and used as the expression centroid.

### Extracellular recording analysis

#### Spike sorting and cell type classification

JRClust^113^ (https://github.com/JaneliaSciComp/JRCLUST) with manual curations was used for spike sorting. We used a combination of quality metrics (described in Majumder et al., 2026^114^) to select single units for analysis. See Extended Data Fig. 5 for the number of recorded neurons. Putative pyramidal and inhibitory neurons were distinguished by spike width (≥ 0.5 ms and <0.35ms respectively^107^). Unless otherwise specified, both cell types were included in the analyses.

Peri-stimulus time histogram (PSTH; Figs. 3c,d, and g, 6g and Extended Data Fig. 8, and 11) was calculated using 10-ms time bins and smoothed with a 200 ms causal boxcar filter. In Fig. 3g, PSTHs were z-scored using the mean and standard deviation of spiking activity during the −3 to - 1 s window relative to the Go cue in non-test trials.

#### Correlation in neural population activity (Fig. 3h, i and Extended Data Fig. 6m)

To plot the correlation in neural population activity, we calculated the mean z-scored spike activity of individual neurons in lick right trials to yield a population activity matrix, with the number of rows equal to the number of neurons and the number of columns equal to the number of time points. Then, we computed the temporal autocorrelation matrix of this population activity matrix by calculating the Pearson correlation between population activity vectors at all pairs of time points. To quantify the transition in population activity around the Go cue, we calculated the mean correlation between time points before and after the Go cue for each correlation matrix.

#### Autocorrelation analysis (Fig. 3j-l)

To evaluate the trial-to-trial autocorrelation of ITI activity, we first calculated the mean firing rate of each neuron during the ITI (−5 to 0 s relative to the Go cue) for every trial. We then computed the autocorrelation of this trial-by-trial activity for each neuron. The autocorrelation at lag 10 was used for the quantification in Fig. 3k and l. In Fig. 3k, time constants were mapped onto a two-dimensional anatomical coordinate system (coronal). Cells were grouped into spatial bins (125 × 125 µm), and the mean time constant was calculated for each bin. The resulting map was spatially smoothed using a 3 × 3 bins median filter. Time constants were represented by color using the parula colormap, while brightness reflected the number of cells within each bin.

#### *Poisson GLM analysis* (Fig. 3e,f and Extended Data Fig. 6)

For each neuron, spiking activity was modeled using a Poisson generalized linear model (GLM):

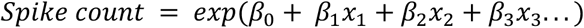

where the regressors included upcoming choice (UC), previous choice (PC), previous reward amount (PA), previous reward delay (PD), the interaction between previous reward amount and previous choice (PA × PC), and the interaction between previous reward delay and previous choice (PD × PC). The interaction terms were calculated after z-scoring the corresponding regressors. Spiking activity during ITI (spike count between −5 to 0 s relative to the Go cue on each trial) or after the Go cue (0-0.5 s) was analyzed. Neurons with less than 300 trials were excluded from analyses.

Models were fit using L2-regularized regression, with the regularization parameter selected by 10-fold cross-validation to minimize deviance. To identify neurons selectively encoding each task variable, we applied recursive feature elimination. Model comparisons were performed using repeated two-fold cross-validation (10 repetitions), with folds stratified across the six combinations of previous choice and reward-delay condition. The contribution of each regressor was quantified by the difference in held-out log likelihood between the full and reduced models, and significance was assessed using a one-sided Wilcoxon signed-rank test across cross-validation folds and repetitions. Neurons were considered to significantly encode a task variable at P < 0.005, followed by Benjamini–Hochberg correction for multiple comparisons. As a shuffle control, trial labels were randomly permuted, disrupting the relationship between neural activity and task variables.

To determine whether the proportion of selective neurons in each brain region exceeded chance levels, we performed hierarchical bootstrapping by resampling sessions and neurons with replacement and tested the null hypothesis that the proportions of selective neurons were the same in the data and shuffle control. Statistical significance was assessed using Benjamini–Hochberg correction (α = 0.05).

To characterize similarity in task-encoding profiles across brain regions (Fig. 3f), we constructed a brain region × regressor matrix containing the proportion of significantly selective neurons. We computed pairwise Spearman’s rank correlations between regions based on this matrix and used the resulting correlation matrix to perform hierarchical clustering with average linkage, using 1−r as the distance metric. Regions were reordered according to the resulting dendrogram, and the reordered correlation matrix was visualized.

To quantify within- and across-group similarity between vPFC and other brain regions (*p*-value in Fig. 3f), we used hierarchical bootstrap samples of the region × regressor matrix. For each bootstrap iteration, we computed Spearman’s correlations between regions and compared the mean pairwise correlation among vPFC regions with that between vPFC and other regions. Statistical significance was assessed from the bootstrap distribution of these within-versus across-group differences.

#### Pseudopopulation analysis (Extended Data Fig. 9h-k)

For pseudopopulation analyses, neurons recorded across sessions were pooled after calculating the mean selectivity for each block type (with different reward-amount ratios or delays). Analyses were restricted to stay trials, because switch trials were rare in the short-delay blocks, allowing fair comparisons across delay and amount ratio conditions. For each neuron and block type, choice selectivity was calculated as the difference in mean z-scored spiking activity between the two lick directions. The similarity between the choice selectivity vectors for two block types was quantified using cosine similarity,

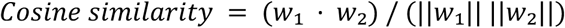

where *w*₁ and *w*₂ denote the choice selectivity vectors for the two block types. A cosine similarity of 1 indicates aligned representations, 0 indicates orthogonal representations. Negative values indicate oppositely oriented representations.

Neurons with at least 7 stay trials per each block type (either no delay and delay, or base ratio and test ratio) per lick direction (left and right) were included. To control for differences in the number of recorded neurons across brain regions, pseudo-population size was matched to the smallest number of eligible neurons across regions. Values were summarized as the mean and SEM across bootstrap iterations (neurons were resampled with replacement).

### Single-Session Population Analyses

For analyses based on single sessions (Figs. 4-6 and Extended Data Figs. 7, 9, and 13), neurons with mean firing rates <0.1 Hz were excluded. Trial ranges were defined to include 70% of neurons, and neurons that did not span the entire trial range were excluded. Following this preprocessing, we included sessions with at least 10 neurons and at least three blocks of each of the six block types (left or right large-reward side with 0-, 0.5-, or >1-s delay) to minimize the impact of slow signal drift on the results. In addition, we performed a circular permutation analysis, as detailed below. Spiking activity was binned in 10-ms time windows. Activity from −5 to 0 s to the Go cue (ITI activity) was analysed on each trial, except for the post-Go-cue activity (from 0 to 0.5 s after the Go cue) analysis in Extended Data Fig. 7o-q.

#### *Participation ratio* (Extended Data Fig. 9l)

For each brain area, we randomly subsampled 15 neurons and calculated the participation ratio of the resulting population activity following principal component analysis (PCA) in neuronal space. The participation ratio was calculated as (Z*λ_i_*)^2^/ Z*λ*^2^, where *λ_i_* are the positive eigenvalues of the covariance matrix. A higher participation ratio indicates higher effective dimensionality of the population activity.

*Decoding and coding direction analysis of upcoming choice (CDupcoming)*

Decoding analyses were performed separately for each recording session. Population activity during the ITI (−5 to 0 s from cue) was used to predict the upcoming choice using a logistic regression classifier with L2 regularization. Z-scored spike rate from simultaneously recorded neurons within the analysis window served as input features, and upcoming lick direction (left versus right) was the binary response variable. The regularization strength was determined by five-fold cross-validation for each session.

To ensure that decoding performance reflected upcoming choice rather than previous choice, equal numbers of stay and switch trials were randomly subsampled before training the decoder. Decoder performance was evaluated using leave-one-out cross-validation, such that each trial was projected using a decoder trained on all other eligible trials, and decoding accuracy was quantified as the area under the receiver operating characteristic curve (AUC). As a control, we performed a circular permutation analysis in which trial labels were circularly shifted by a random offset for 1,000 iterations. This preserved the trial-to-trial autocorrelation while disrupting the relationship between neural activity and choice.

We defined the coding direction for upcoming choice **CDupcoming** as the weight vector (**w**) of the fitted logistic regression decoder. To quantify trial-by-trial choice representations, we computed the decoder logit (linear predictor) for each held-out trial using its corresponding leave-one-out decoder and population activity at each time point within the trial:

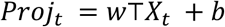

Where *X_t_* denotes the neural population activity (250-ms boxcar filtering with a 50-ms step size) and *b* is the intercept of the fitted logistic regression model. This scalar value (decoder logit) is zero at the decision boundary, corresponding to a predicted choice probability of 0.5, with positive and negative values indicating evidence favoring opposite choices. The resulting scalar value represented neural activity along **CDupcoming** and was used for subsequent analyses.

Similarly, for each recording session, separate L2-regularized logistic regression classifiers were trained to discriminate upcoming lick direction using only stay trials from blocks with different reward delays (>1-s, 0.5-s, and 0-s). Stay trials were used because switch trials were rare in the short-delay blocks, allowing fair comparisons across delay conditions. We defined the coding directions for the >1-s delay (**CDdelay**) and 0-s delay (**CDno delay**) blocks as the weight vector of the fitted logistic regression decoders. Decoder logits were computed using the same leave-one-out procedure described above.

For optogenetics (Fig. 6), population activity during the 500-ms period preceding the decision (−0.5 to 0 s) was used to maximize decoding performance during the stimulation period. Analyses were restricted to sessions with reliable choice decoding (10-fold cross-validated AUC > 0.8) in control (non-stimulation) trials. To pool sessions with different reward contingencies, **CDupcoming** was sign-aligned to the large-reward side, such that positive values always indicated a decoded bias toward the large-reward option.

To determine whether vPFC silencing differentially altered choice representations in delay and no-delay blocks, mean **CDupcoming** was calculated before photostimulation (−1.2 to −0.7 s) and during photostimulation (−0.5 to 0 s). Statistical significance was assessed using a permutation test in which delay and no-delay block labels were randomly shuffled within each session while preserving trial numbers (1000 iterations). To account for animal-to-animal variability, sessions were resampled with replacement in every permutation iteration.

### Statistics

The sample sizes were similar to the sample sizes used in the field. No statistical methods were used to determine the sample size. During spike sorting, experimenters could not tell the trial type and, therefore, were blind to conditions. All signed-rank and rank-sum tests were two sided. All bootstrapping and permutation tests were performed over 1,000 iterations.

## Data availability

The recording data in NWB format and codes will be made publicly available through the DANDI Archive and GitHub upon publication.

## Extended Data Figure

**Extended Data Figure 1.**
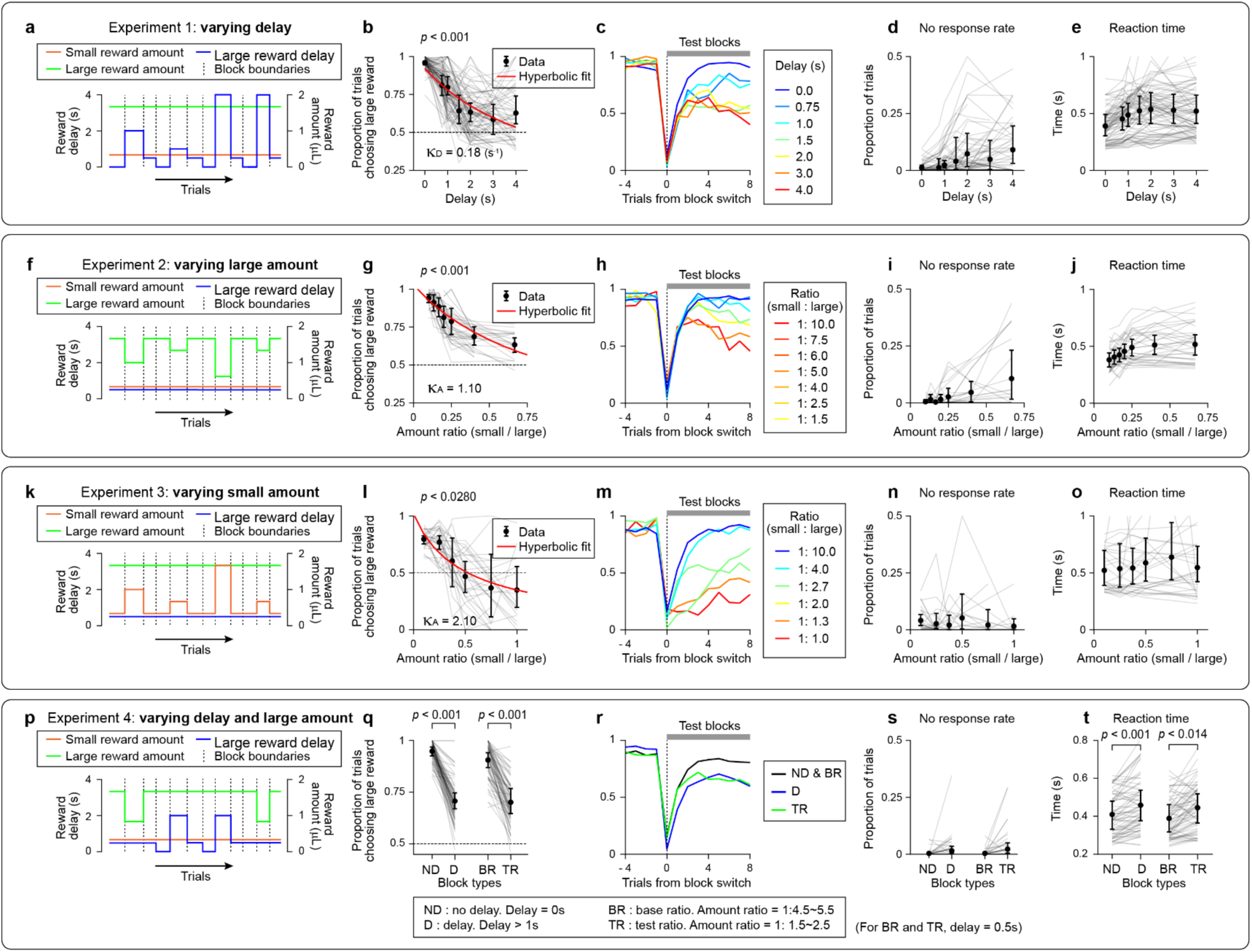
Summary of behavioral effects of reward amount and delay manipulations. Behavioral characterization across different test conditions: Varying reward delay (**a-e**, n = 7 mice); Varying large-reward amount (**f-j**, n = 7 mice); Varying small-reward amount (**k-o**, n = 5 mice); Varying reward delay and amount in the same session (**p-t**, n = 6 mice). Varying reward delay or reward amount independently in separate sessions, or varying both within the same session, produced similar effects. Increasing the small-reward amount delayed switching choice after block switch (**o**), indicating that the relative value of the small reward had increased. **a,** Example session in which reward delay was randomly varied across interleaved test blocks. The small and large reward amounts were held constant. **b,** Proportion of trials choosing the large reward as reward delay varied. Line, individual session. Circle, mean; bar, 95% confidence interval (CI; hierarchical bootstrap); red line, hyperbolic fit; κ_D_, temporal discount coefficient (Methods). **c,** Proportion of trials choosing the large reward aligned to block switches for test blocks with different reward delays. Color, delay duration. **d,** Proportion of trials with no response as reward delay varied. Same format as in **b**. **e,** Reaction time as reward delay varied. Only trials in which the upcoming choice was the large-reward option were included. Same format as in **b**. **f-j,** same as in **a-e** for varying large-reward amount. Reward delay was held constant at 0 s for both small and large reward options. **k-o,** same as in **a-e** for varying small-reward amount. Reward delay was held constant at 0 s for both small and large reward options. **p-t,** same as in **a-e** for varying both reward delay and amount.

**Extended Data Figure 2.**
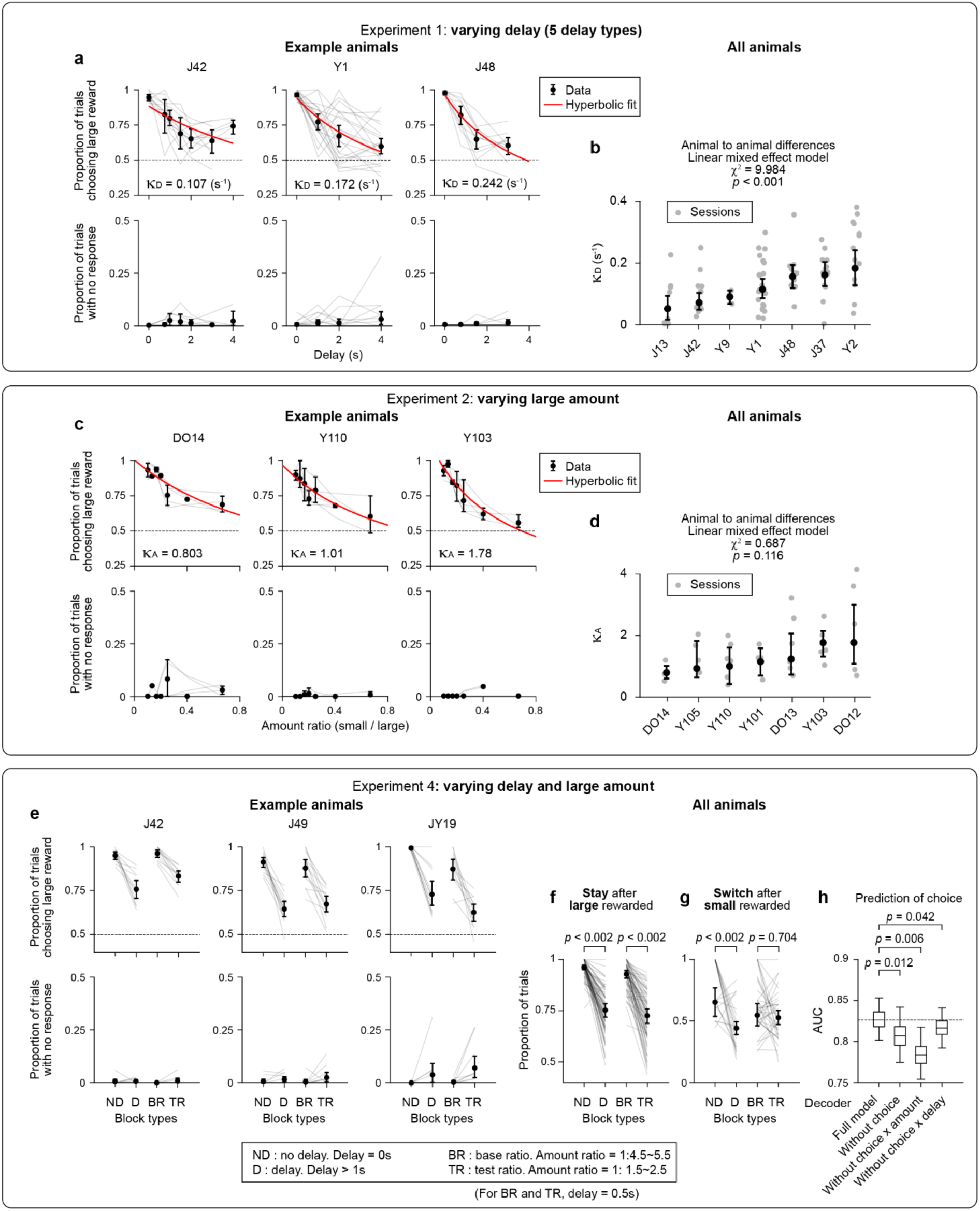
Behavior of individual animals. Behavior of individual animals. The extent of temporal discounting varied across animals. **a,** Top, proportion of trials choosing the large reward as reward delay varied in example mice. Line, individual session. Circle, mean; bar, 95% confidence interval (CI; hierarchical bootstrap); red line, hyperbolic fit; κ_D_, temporal discount coefficient (Methods); Bottom, proportion of trials with no response as reward delay varied. **b,** Distribution of discounting rates across animals. Gray circles, individual sessions; Black, mean ± 95% CI for each animal. Significant differences in discounting rates were observed across animals (linear mixed effect model), indicating stable individual variability in temporal discounting behavior. **c-d,** Same as **a-b** for varying large-reward amount. **e,** Same as **a** for varying both reward delay and amount (instead of delay duration, performance across different block types is shown). **f,** Proportion of trials in which animals stayed licking in the same direction after receiving a large reward (win-stay), across different blocks. *p*-values, hierarchical bootstrap test comparing means with *Bonferroni* correction for two comparisons. n = 6 mice (reward amount and delay were varied within the same session to enable within-animal comparisons: **f-h**). **g,** same as in **f** for proportion of trials in which animals switched licking direction after receiving a small reward (lose-switch), across different blocks. **h,** Performance of the regression model in Fig. 2g compared with reduced models (AUC, two-fold cross-validation), confirming that all regressors contribute to predicting upcoming choice. *P*-value, hierarchical bootstrap.

**Extended Data Figure 3.**
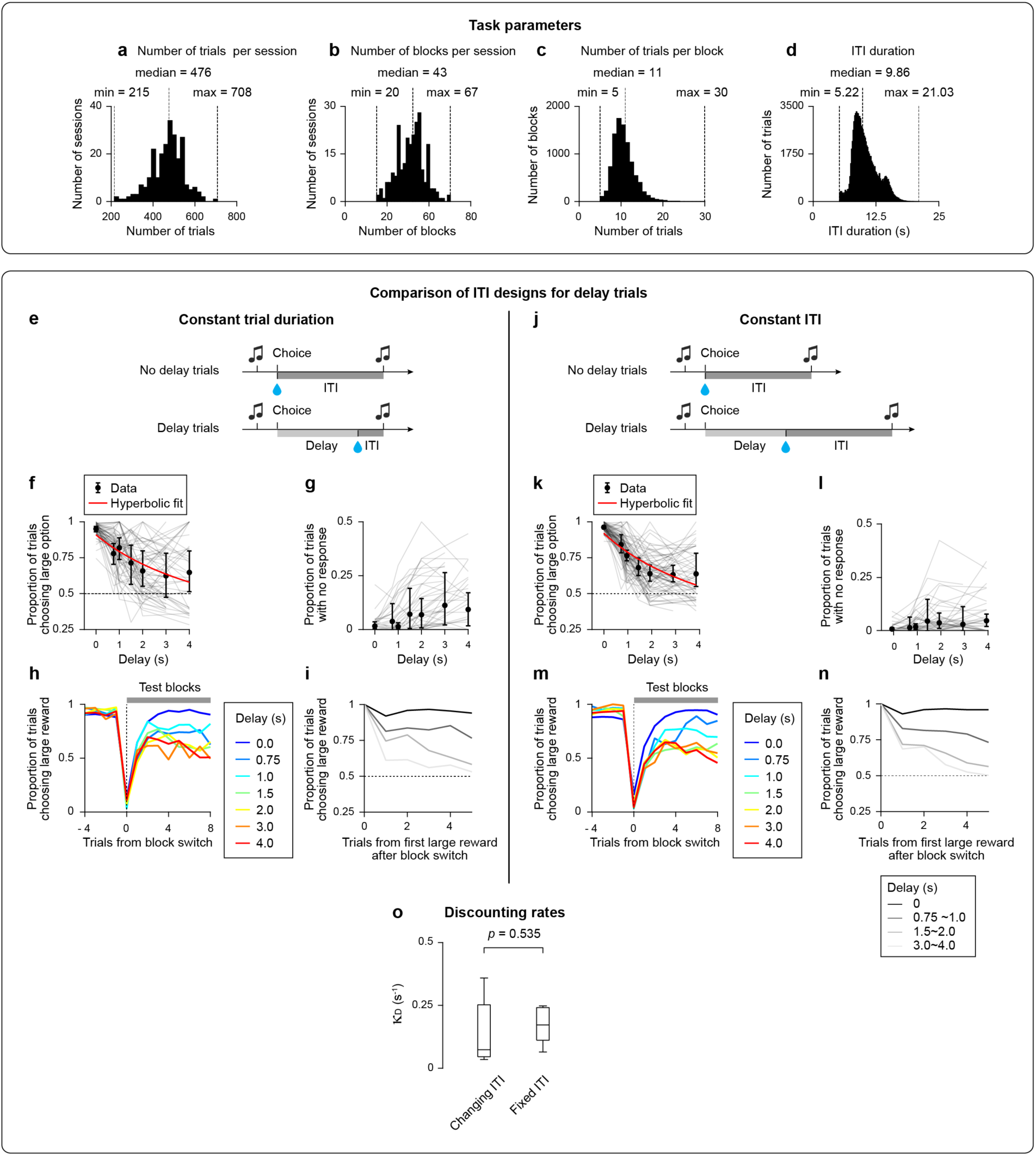
Task parameters and comparison of inter-trial interval (ITI) designs in the temporal discounting task. An alternative account posits that animals maximize overall reward rate rather than discounting reward delay^44^. We therefore compared two task variants: (i) task duration equalized, where longer delays were followed by shorter ITIs to equalize trial durations across trials with distinct delays (**constant trial duration; e-i**, n = 7 mice); and (ii) task duration varies due to constant ITI duration (**constant ITI; j-n**, n = 7 mice). Discount rates did not differ between variants (**o**; *p* = 0.535, *p* value, two-sided Wilcoxon rank-sum test), thus reward rate maximization alone cannot explain the behavior, indicating that mice exhibit genuine temporal discounting in this task. For all other experiments we used the constant ITI version to keep ITI durations comparable across trials. **a,** Distribution of the number of trials per session. **b,** Distribution of the number of blocks per session. **c,** Distribution of the number of trials per block. **d,** Distribution of ITI durations. Median, minimum, and maximum values are indicated. **e,** Schematic of the constant trial duration design. **f,** Proportion of trials choosing the large reward as reward delay varied (reward magnitude was held constant). Line, individual session. Circle, mean; bar, 95% confidence interval (CI; hierarchical bootstrap); red line, hyperbolic fit (Methods). **g,** Proportion of trials with no response as reward delay varied. Same format as in **f**. **h,** Proportion of trials choosing the large reward aligned to block switches for test blocks with different reward delays. Line, grand mean across mice. Color, delay duration. **i,** Proportion of trials choosing the large reward aligned to the first large-reward trial after a block switch. The probability of switching away from the large-reward side depended on the reward delay associated with the large reward. **j-n,** Same as in **e-i**, but for the **constant ITI** design. **o,** Comparison of κ_D_, temporal discount coefficient, between the constant trial duration and ITI task designs. No significant difference was observed between the two conditions.

**Extended Data Figure 4.**
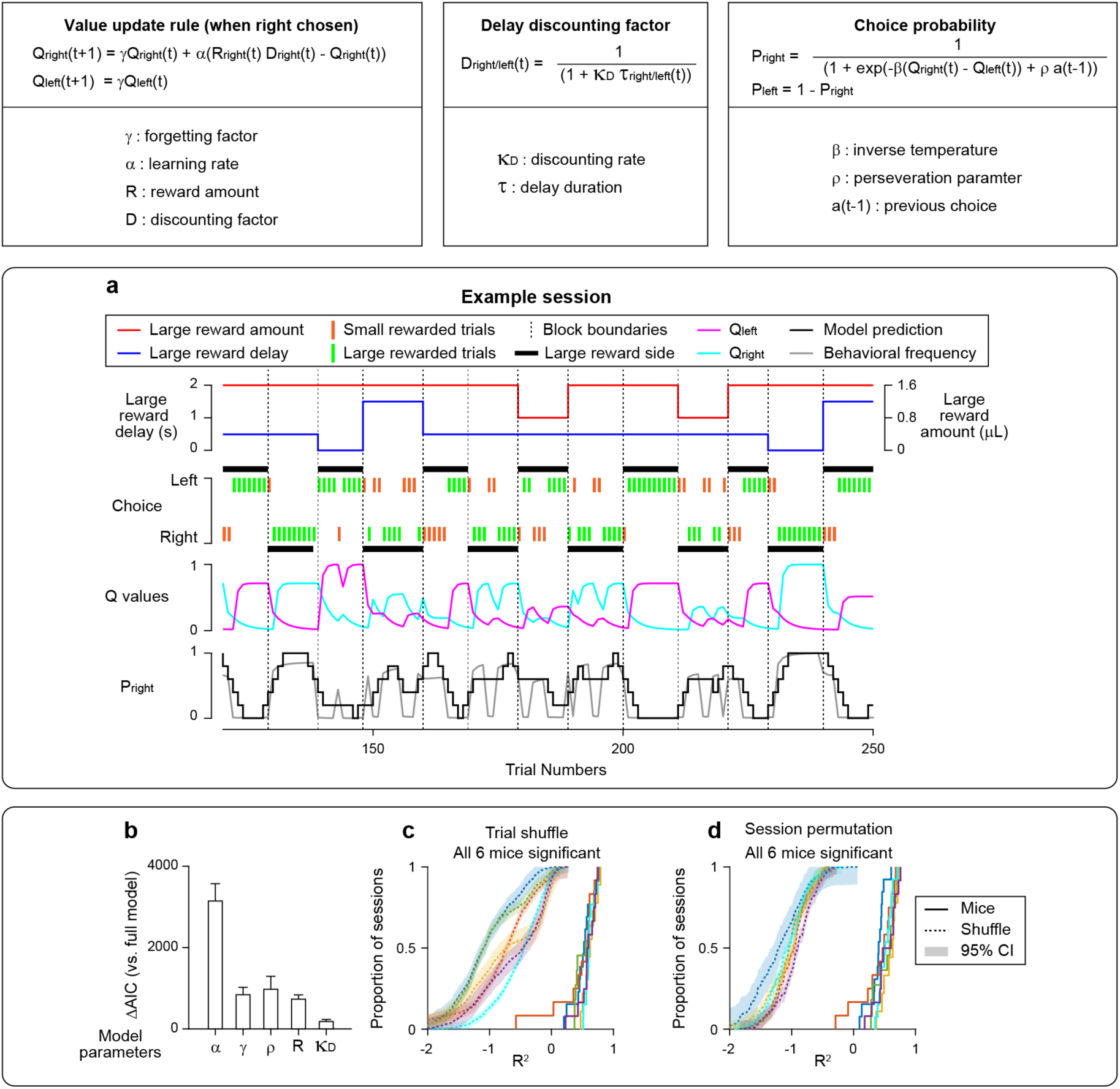
Q-learning model fit. **a,** Example behavioral session and Q-learning model fit. Top, reward delay (s; blue) and reward amount (µl; red) of the large-reward choice. Second row, trial-by-trial choices and reward outcomes under the block schedule (dashed lines and black boxes). Third row, estimated action values (Q-values) for the left and right choices. Fourth row, model-predicted probability of choosing the right option (black) and the observed choice behavior (gray; boxcar-smoothed over 5 trials). **b,** Contribution of individual model parameters to model fit assessed using the Akaike Information Criterion (AIC), evaluated by removing it from the full model and quantifying the resulting change in model fit. **c-d**, Model performance on held-out trials exceeded that of the trial-shuffle control (trial types shuffled within sessions; **c**) and the session-permutation control (trial types swapped across sessions; **d**), as evaluated by *R*² between model-prediction and observed choice behavior (gray vs. black line, fourth row of **a**).

**Extended Data Figure 5.**
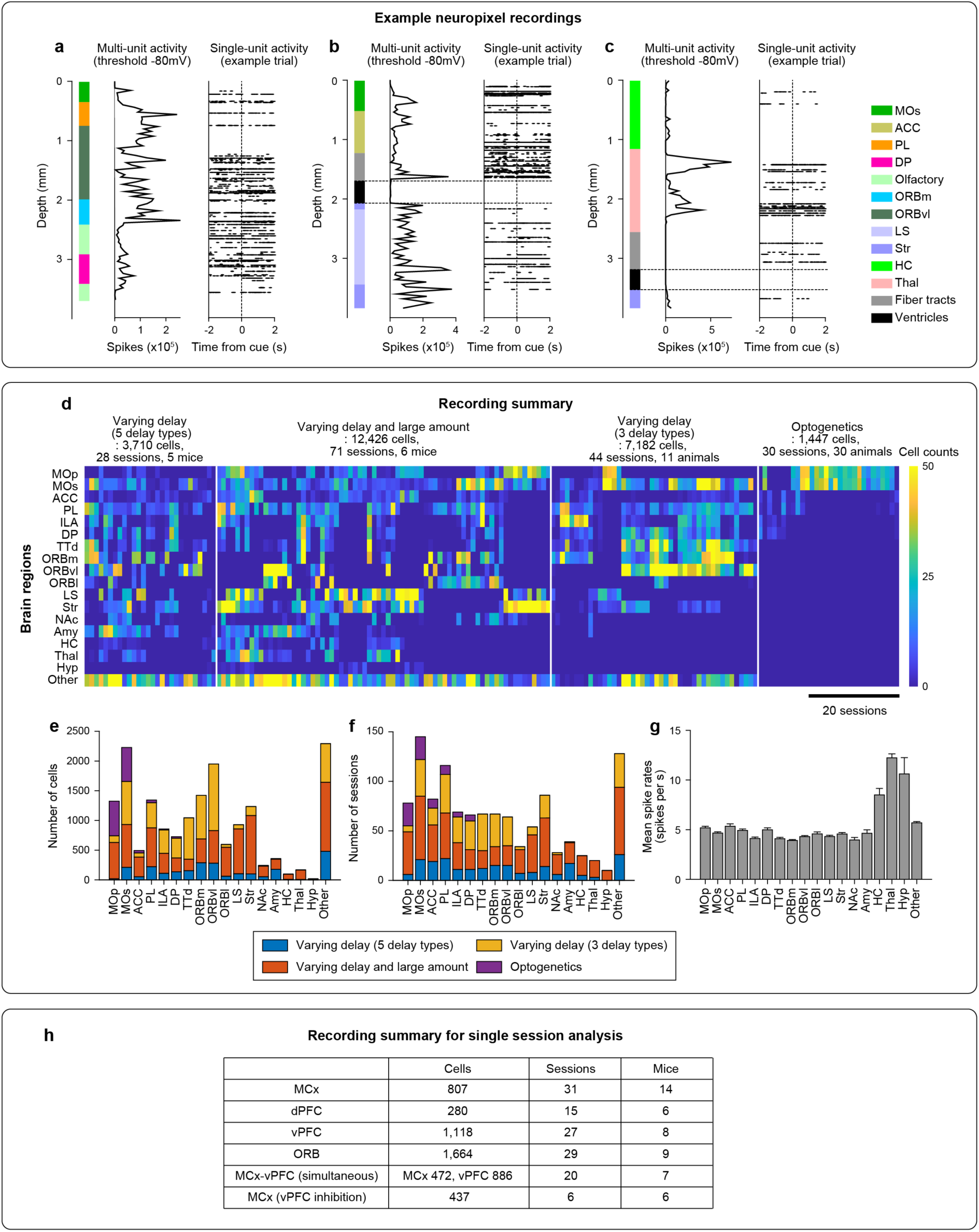
Recording quality and distribution of units. **a-c,** Example Neuropixels recordings. Left, inferred recording sites registered to Allen CCF. Color, brain regions traversed by the probe. Middle, multi-unit activity (threshold = −80 μV) as a function of recording site. Right, example single-unit activity in an example trial. Note the absence of spiking activity in the ventricles (black box), validating the accuracy of the registration. **d,** Distribution of recorded units across sessions and brain regions. Heat map, the number of isolated units for each recording session, organized by brain region and experiment type. **e,** Total number of recorded units across all sessions for each brain region. Stacked bars indicate contributions from different experiments. **f,** Number of recording sessions containing at least one isolated unit in each brain region. Colors indicate experiment type. **g,** Grand mean firing rate across all recorded units for each brain region during ITI. Bars represent mean ± SEM across units. Brain regions include M1, M2, ACC, PL, IL, DP/TTd, mOFC, vOFC, lOFC, LS, striatum (Str), nucleus accumbens (NAc), amygdala (Amy), hippocampus (HC), thalamus (Thal), and hypothalamus (Hyp). **h,** number of neurons, sessions, and animals analyzed for single-session analysis (Fig. 4-6) and simultaneous recording (Extended Data Fig. 9l-n).

**Extended Data Figure 6.**
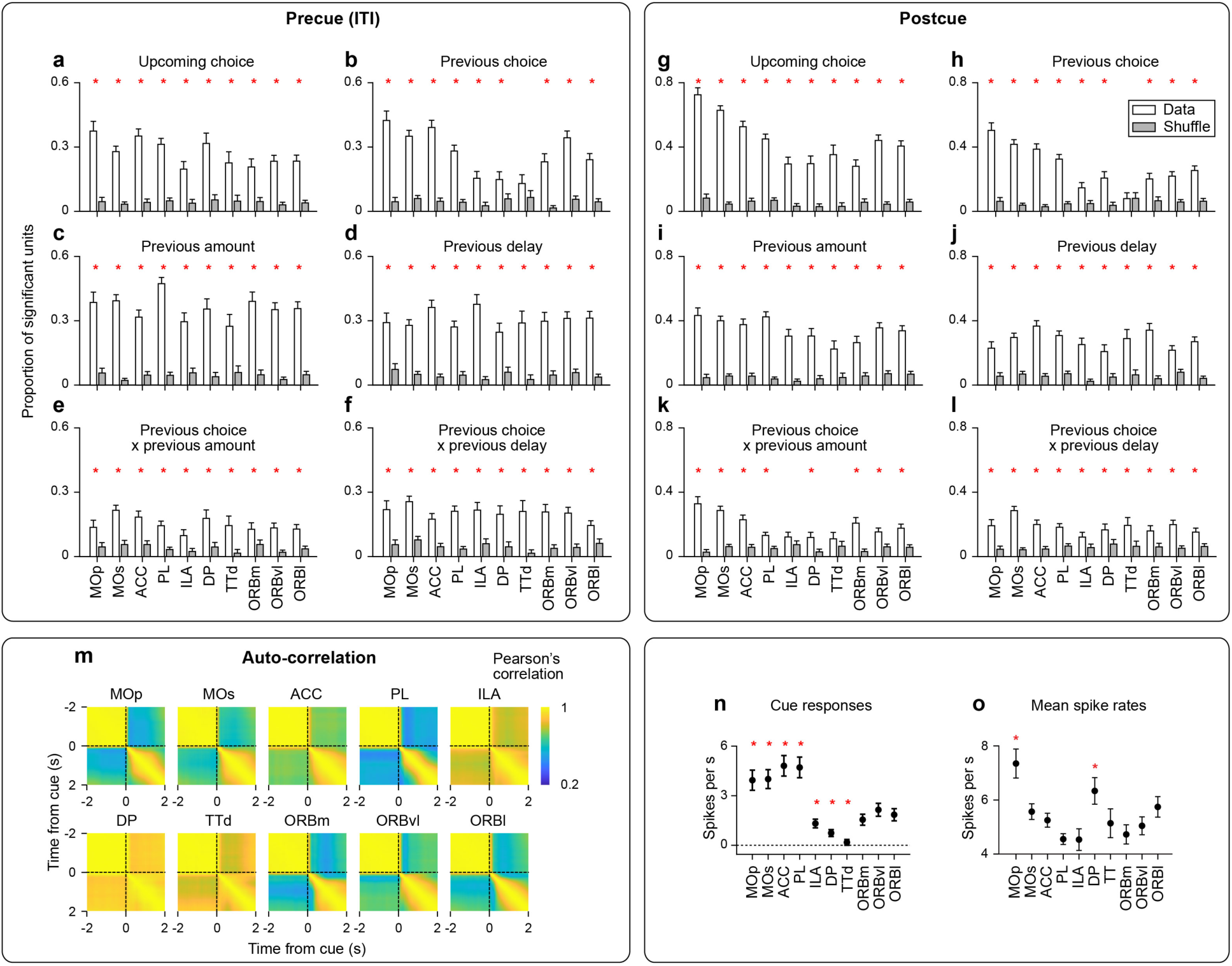
Regression analysis of spiking activity across brain areas. **a-f,** Proportion of cells significantly encoding each task variable during ITI (−5 - 0 s from the go cue) in each brain area. Box, mean; bar, SEM (hierarchical bootstrap); *, *p*< 0.05 Hierarchical Bootstrap testing the null hypothesis that proportion of significant units is the same between data and shuffle controls, followed by Benjamini–Hochberg procedure. **g-l,** Proportion of cells significantly encoding each task variable during post cue period (0-0.5 s from the go cue) in each brain area. Same format as in **a-f**. **m,** Pearson’s correlation matrix comparing population activity across time points in each area. Same format as in Fig. 3h. **n,** Cue-evoked responses across brain areas, quantified as the change in firing rate from the pre-cue (ITI) to post-cue period. Circle, mean; bar, SEM (hierarchical bootstrap). *, *p* < 0.05, leave-one-region-out bootstrap test comparing each region with the mean of the remaining regions. **o,** Mean spike rate during the ITI (−5 - 0 s from the go cue) in each brain area. Same format as in **n**. This metric quantifies how many of the six task variables contribute to the neural representation. The participation ratio is high (∼5), indicating that task encoding is distributed across multiple task variables (heterogeneous, combinatorial coding) rather than concentrated on a few (categorical coding)^60^. Moreover, it does not differ significantly across brain areas (Kruskal-Wallis test).

**Extended Data Figure 7.**
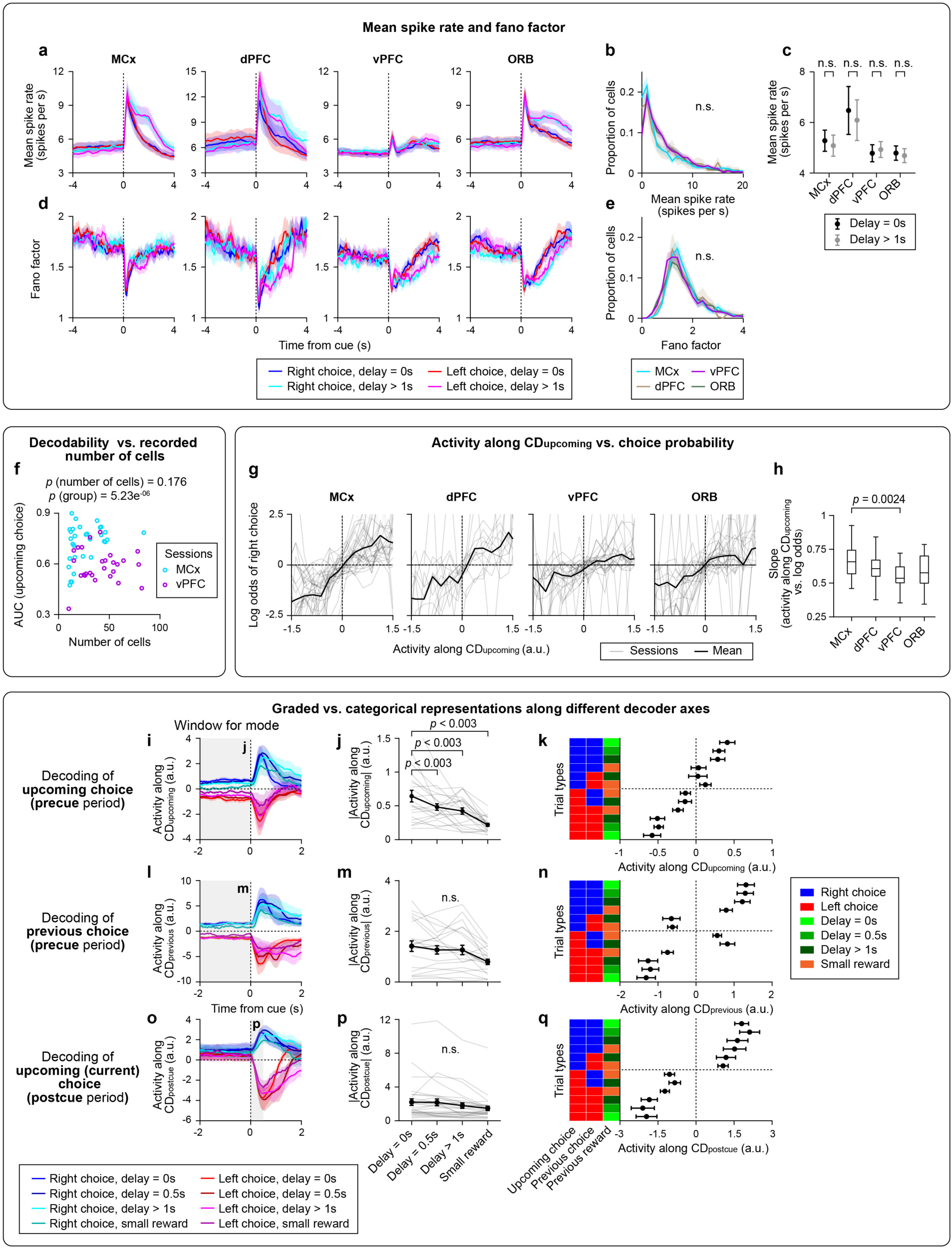
Summary of single-session activity. Basic firing properties (firing rate and Fano factor) were similar across brain regions (**a–e**). MCx ITI activity projected along **CD_upcoming_** exhibited graded activity patterns according to previous trial type (**i-k**), whereas ITI activity projected along **CD_post cue_** or **CD_previous_** exhibited more discrete representations of previous choice (**l-q**). **a,** Grand mean spike rate in each brain area. Line, mean; shade, SEM. n = 31, 15, 27, and 29 sessions in MCx, dPFC, vPFC, and ORB, respectively. **b,** Distribution of mean spike rate during ITI (−5 - 0 s before the go cue) across cells in each area. Line, mean; shade, SEM. n.s., bootstrap test of the null hypothesis that mean spike rates were equal between each pair of brain regions (p > 0.05 for all pairwise comparisons). **c,** Comparison of grand mean ITI activity between delay = 0s (no-delay) and delay > 1s (delay) blocks. n.s., p > 0.05 for bootstrap test of the null hypothesis that grand mean spike rates were equal between blocks. **d-e,** Same as **a-b** for Fano factor. **f,** Decoding of upcoming choice in MCx (cyan) and vPFC (purple) as a function of the number of recorded units. Even after accounting for the number of recorded units, MCx showed higher decodability than vPFC (*p*-values, linear mixed-effects model). **g,** Population activity in each area projected along **CD_upcoming_** vs. log odds of a rightward choice. Activity was binned every 0.25-a.u. to calculate the log odds. Thick line, mean; thin line, session. **h,** Session-wise linear regression slope between **CD_upcoming_** activity and the log odds of a rightward choice (**g**) in each brain area. *p*-value, hierarchical bootstrap comparing means with *Bonferroni* correction for six pairwise comparisons following a Kruskal-Wallis test (p = 0.00672). All other pairs were *p*> 0.05. **i,** MCx population activity projected along **CD_upcoming_**. Color, trial type. Line, mean; shade, SEM. Reproduced from Fig. 4c for comparison. **j,** Absolute value of **CD_upcoming_** activity as a function of previous trial types. *P*-value, bootstrap test comparing the mean for Delay = 0s vs. other conditions followed by *Bonferroni* correction for three comparisons. Reproduced from Fig. 4d for comparison. **k,** MCx **CD_upcoming_** activity across trial types. Circle, mean; bar, SEM. **l-n**, Same as **i-k** for activity along **CD_previous_** (decoder of previous choice based on ITI activity; Methods). **m,** *p* = 0.0795, Kruskal-Wallis test. **o-q,** Same as **i-k** for activity along **CD_post cue_** (decoder of choice based on post-cue activity; Methods). **p,** *p* = 0.350, Kruskal-Wallis test.

**Extended Data Figure 8.**
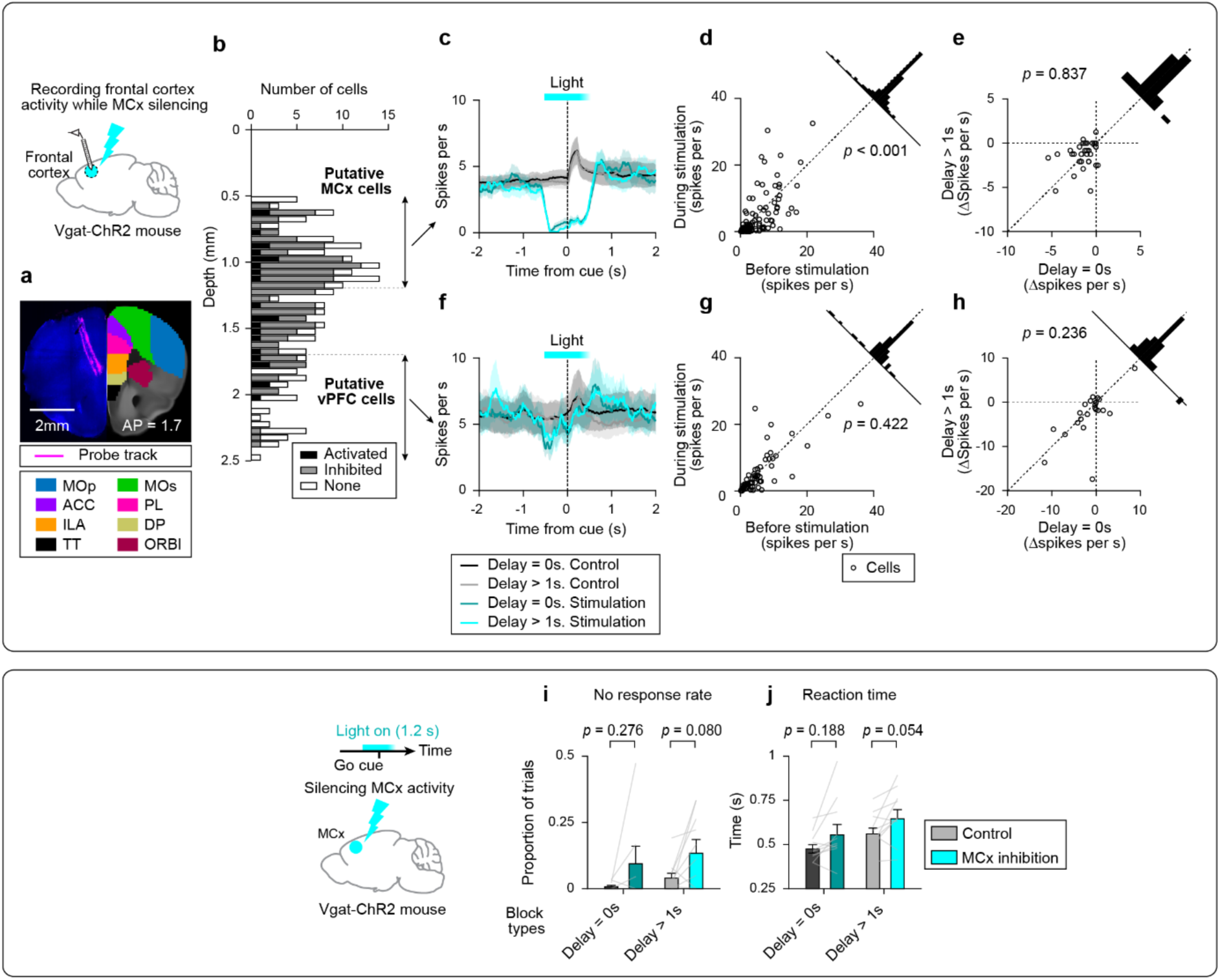
Validation of MCx optogenetic inhibition. Validation of MCx inhibition **(a-h)** and its behavioral effect **(i-k). a,** Example brain for MCx recording during bilateral MCx silencing in Vgat-ChR2 mouse. **b,** Distribution of recorded cells across depth. Color, cells classified as activated, inhibited, or unaffected by MCx silencing (signed-rank test, ⍺ = 0.05, n = 313 neurons). **c,** Grand mean PSTH of putative MCx cells. Gray, control; cyan, MCx silencing. Shading, SEM. Cyan bar, light stimulation. **d,** Spike rate of putative pyramidal MCx neurons (depth < 1.2 mm) before (−0.6-0 s) vs. during (0-0.6 s) the stimulation. Top right, histogram of spike rates during minus before stimulation. *p-*value, signed-rank test between spike rate before and during stimulation. **e,** Change in spike rate during MCx inhibition in delay vs. no-delay blocks. The same format as in **d**. **f-h,** Same as **c-e** for putative vPFC cells (depth > 1.8 mm). **i-j,** No response rate and reaction time during MCx silencing. Same format as in Fig. 4j (n = 10 mice).

**Extended Data Figure 9.**
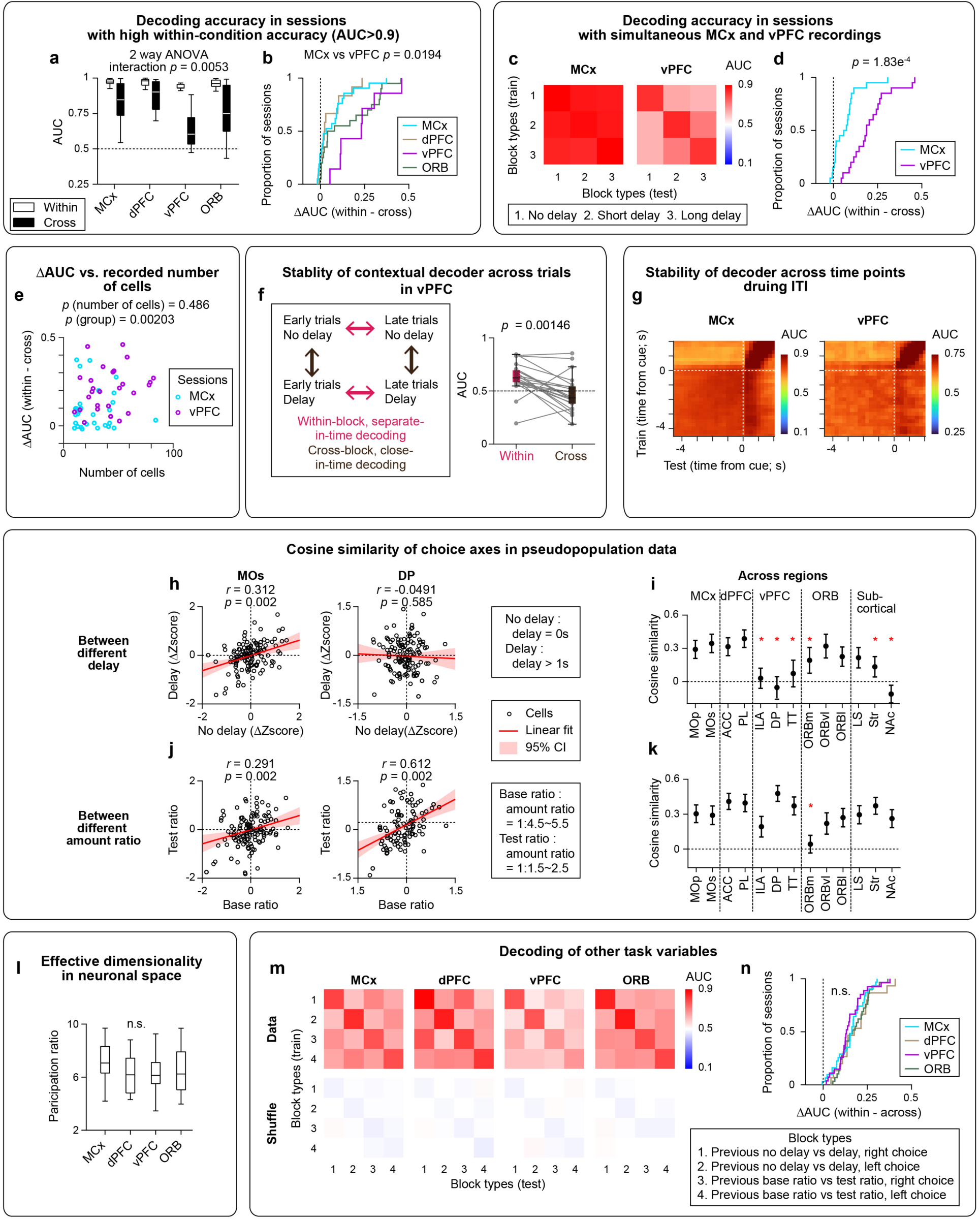
Generalization of population representations across task conditions. **a,** Session-wise decodability, among sessions in which within-block AUC >0.9. In boxplots, the central line, median; box edges, the 25th and 75th percentiles; whiskers, the most extreme values within 1.5× the interquartile range. **b,** Cumulative distribution of the difference between within-block and cross-block decodability across sessions, among sessions in which within-block AUC >0.9. Same format as in Fig. 5d. **c,** Decoding analysis in MCx and vPFC in simultaneous recording sessions (n = 20 sessions). Same format as in Fig. 5b. **d,** Same format as in **b** for simultaneous recording sessions. *P*-value, Kolmogorov–Smirnov test. **e,** ΔAUC (difference between within-block and cross-block decodability) in MCx (cyan) and vPFC (purple) as a function of the number of recorded units (*p*-values, linear mixed-effects model). **f,** Apparent context-dependent coding in vPFC could arise from the temporal proximity of training and test trials in within-condition decoding, particularly given the long trial-to-trial timescale of vPFC activity. To test this possibility, in sessions with sufficient trials (>20 stay trials for each lick direction in both no-delay and delay conditions), we trained separate decoders on the first or last 10 trials of each condition. Each decoder was tested on temporally distant trials from the same condition and temporally proximal trials from the other condition. Decoders performed significantly better in the ‘within-block, separated-in-time’ condition than in the ‘cross-block, close-in-time’ condition for both no-delay and delay decoders. Thus, context-dependent decoding persisted across temporally separated trials. **g,** Stability of the lick-direction decoder across time during the ITI. A decoder trained in the no-delay block, stay-trials using a 500-ms bin and 250-ms sliding window was tested across time points. Decoder performance remained stable throughout the ITI, indicating no rotation of the coding direction, and changed after the cue. **h,** Correlation of single-neuron choice selectivity between different delay (delay = 0s versus delay > 1s; stay trials only) blocks in MOs (left) and DP (right). Circle, neuron; red line, linear fit; shading, 95% CI; *r*, Pearson’s correlation coefficient. **i,** Similarity of single-neuron choice selectivity quantified as cosine similarity. *, p<0.05 bootstrap test of the null hypothesis that cosine similarity is greater than zero. **j-k,** same as **h-i** for choice selectivity between different amount ratios (amount ratio = 1:1.5 - 2.5 versus 1:4.5 - 5.5). **l,** Participation ratio (effective dimensionality) of PCA eigenvalues from population neural activity in individual sessions (Neuronal space; Methods). It was not significantly different across brain areas (*p* = 0.274, Kruskal-Wallis test). Boxplot, same format as in **a**. **m,** Same format as in Fig. 5b for reward delay (and reward amount) decoders for each choice (e.g., a decoder distinguishing whether the left choice had a reward delay or not). **n,** Same format as in Fig. 5d for reward delay (and reward amount) decoders in **m**. There was no significant difference across brain areas (n.s., *p*>0.05, Kolmogorov–Smirnov test for all six pairwise comparisons).

**Extended Data Figure 10.**
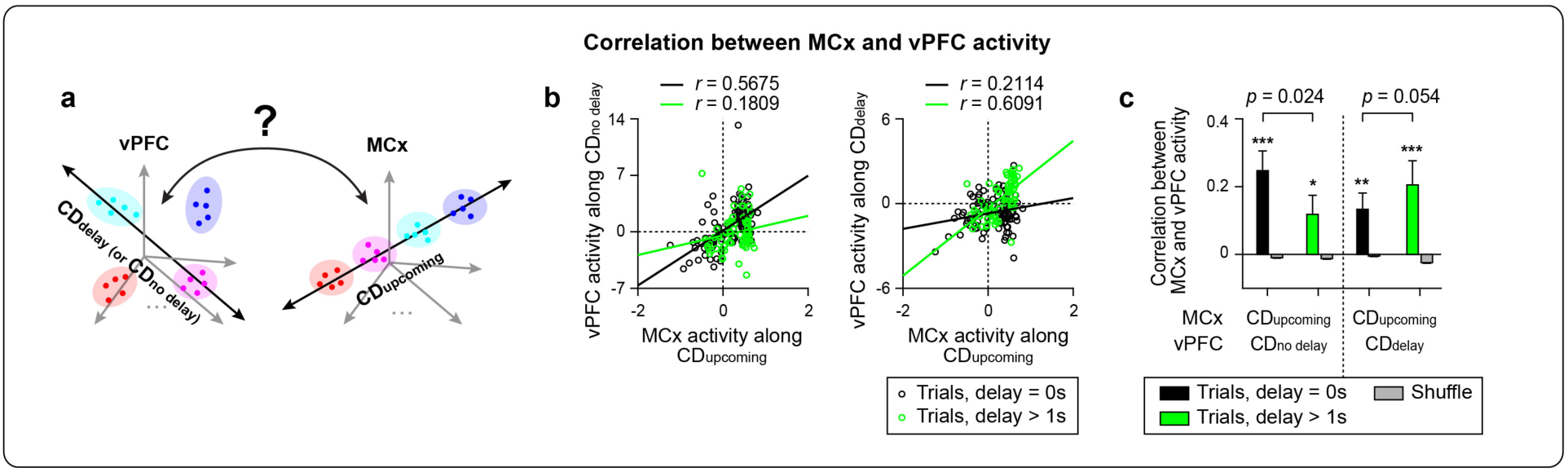
Context-dependent coupling between vPFC and MCx choice activity. **a,** Schematic of the cross-region correlation analysis. **b,** Example session. Relationship between MCx activity along **CD_upcoming_** and vPFC activity along either **CD_no delay_** (left) or **CD_delay_** (right) in different block types (delay or no-delay). Circle, trial. **c,** Trial-by-trial Pearson’s correlation between vPFC activity along **CD_delay_** or **CD_no delay_** and MCx activity along **CD_upcoming_**. Bar, mean; error bar, SEM (bootstrap). Gray, circular permutation. *P*-value, Permutation test of the null hypothesis that the difference in correlation between trial types (delay = 0-s vs. >1-s) is equal to that expected under circular permutation. n = 20 sessions.

**Extended Data Figure 11.**
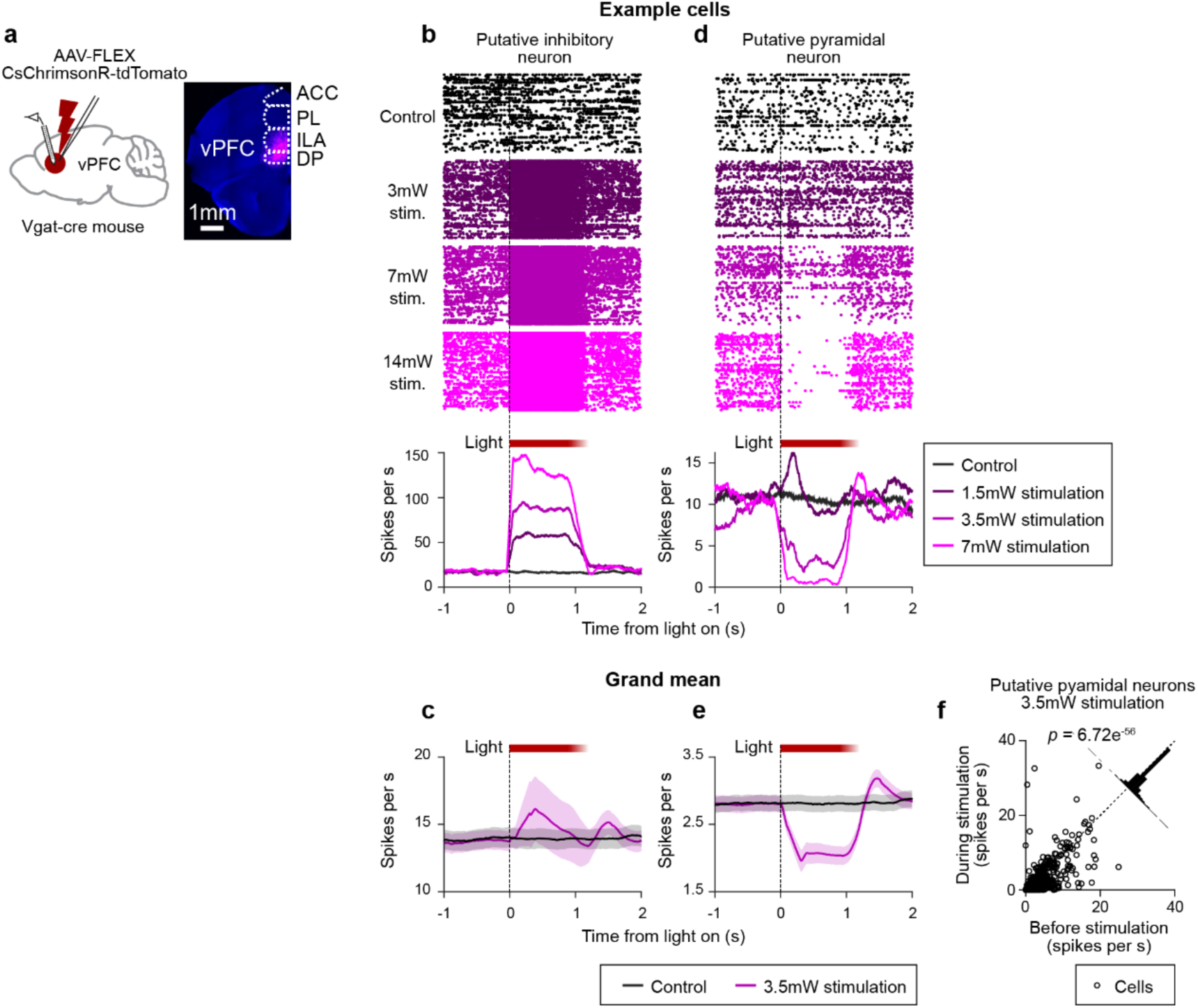
Validation of vPFC optogenetic inhibition. Spiking activity of putative inhibitory neurons increased, whereas that of putative pyramidal neurons decreased, during vPFC photoinhibition, validating the effectiveness of the photoinhibition. **a,** Schematic of recordings in vPFC during vPFC photoinhibition in vGAT-ires-Cre mice expressing ChrimsonR-tdTomato (left). Coronal section from an example animal (right). Blue, autofluorescence; magenta, viral expression (tdTomato). **b,** Spiking activity of an example fast-spiking (putative inhibitory) neuron. Top, spike raster (dots, spikes; rows, trials). Bottom, peri-stimulus time histogram (PSTH). **c,** Grand mean PSTH across putative inhibitory neurons (n = 166 neurons). Shade, SEM (neurons). **d,e,** Same as in **b,c**, respectively, for putative pyramidal neurons (n = 843 neurons). **f,** Spike rate of putative pyramidal neurons before (−0.6∼0 s) vs. during (0∼0.6 s) the stimulation. Top right, histogram of spike rates during minus before stimulation. *p-*value, signed-rank test between spike rate before and during stimulation.

**Extended Data Figure 12.**
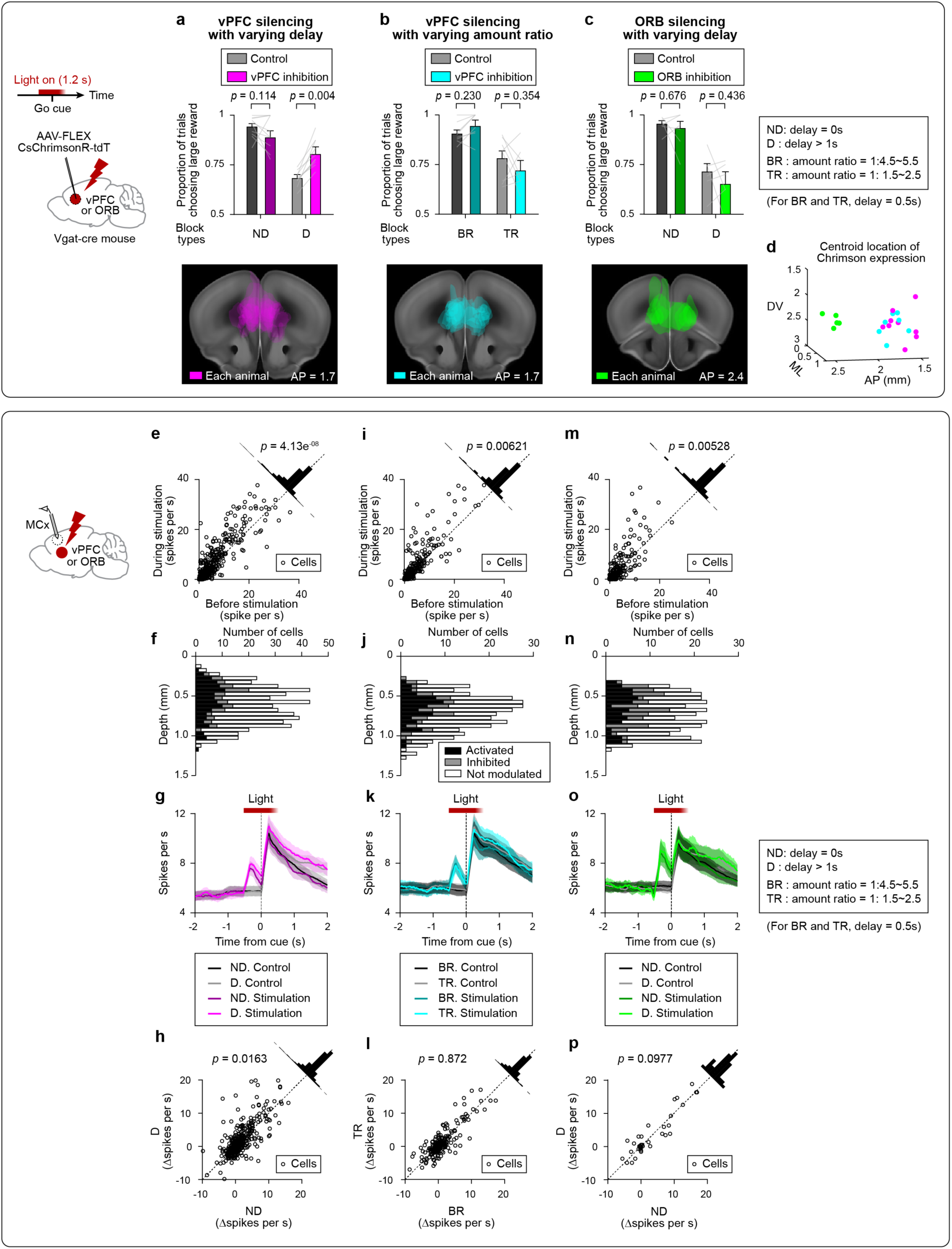
Effects of vPFC and ORB silencing across task conditions. **a,** Top, Proportion of trials choosing the large reward during varying reward delays with vPFC inhibition (n = 10 mice). Bottom, Overlay of virus expression across mice. Reproduced from Fig. 6c**,d** for comparison. **b,c,** Same as in **a** for vPFC inhibition during varying reward amounts (**b**, *n* = 8 mice) and ORB inhibition during varying reward delays (**c**, *n* = 6 mice). **d,** Centroid location of ChrimsonR-TdTomato expression. Circle, mice; color, experiment type. Note that although ORB and vPFC appear similar in coronal sections, they are separated along the AP axis. **e,** Spike rates of MCx putative pyramidal neurons before and during vPFC inhibition. Top right, histogram of the change in spike rate (during minus before stimulation). P value, signed-rank test. Reproduced from Fig. 6f for comparison. **f,** Modulation of spiking activity as a function of recording depth. Activated and inhibited neurons were defined by a signed-rank test comparing before vs. during stimulation (α = 0.05). **g,** Grand mean PSTHs of MCx putative pyramidal neurons during control and stimulation trials. Line, mean; shaded area, SEM (neurons). **h,** Change in spike rate during vPFC inhibition in delay and no-delay blocks. Reproduced from Fig. 6h for comparison. **i-p**, Same as **e-h** for vPFC inhibition during varying reward amounts (**i-l**) and ORB inhibition during varying reward delays (**m-p**).

**Extended Data Figure 13.**
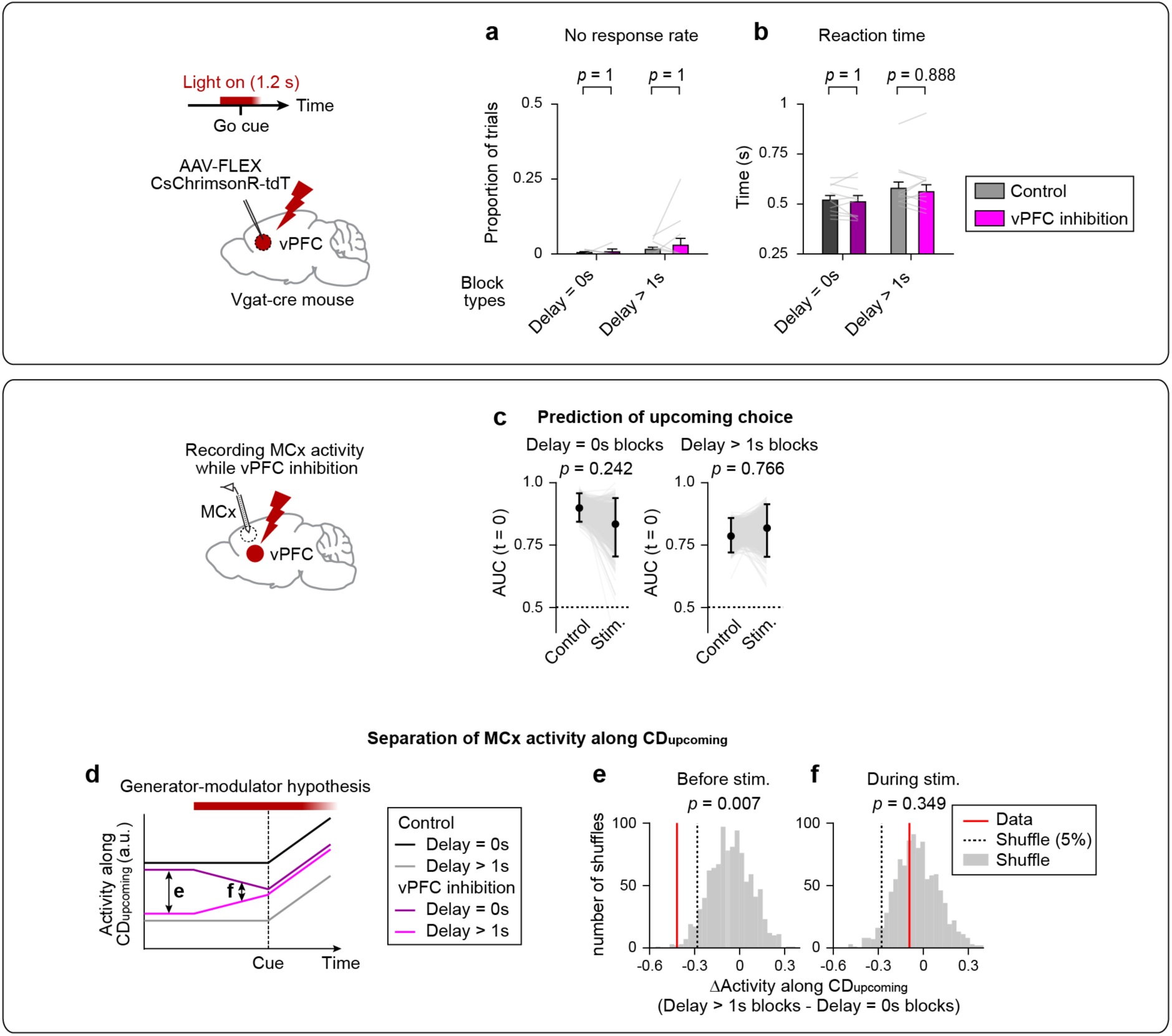
Transient optogenetic perturbation of vPFC. **a-b,** Reaction time and no response rate during vPFC inhibition (n = 10 mice). Same format as in **Extended Data** Fig. 8i**-j**. **c,** Decodability (AUC) of upcoming choice from MCx population activity during vPFC inhibition. A single decoder was trained on control trials and its performance at the go cue (t = 0) was evaluated on stimulation trials. Circle, mean; error bar, 95% CI (bootstrap). *P*-value, bootstrap comparing control vs. stimulation. **d,** Schematic illustrating the predicted effect of vPFC inhibition on context-dependent separation of MCx activity under generator-modulator hypothesis. **e-f,** Effect of vPFC silencing on context-dependent separation of MCx activity along **CD_upcoming_** before (**e**) and during (**f**) stimulation. Red line, data; gray, circular permutation distribution (1000 iterations); gray dotted line, 5th percentile of the permutation distribution. *P*-values, Permutation test with the null hypothesis that the separation in activity is smaller in control.

**Extended Data Table 1.** Summary of experimental datasets.

| Experiments | Relevant figures | Number of animals | Number of sessions | Large reward delay (s) | Small reward amount (μL) | Large reward amount (μL) |
| --- | --- | --- | --- | --- | --- | --- |
| Varying delay<br>(5 delay types,<br>constant trial duration) | Fig. 2e,f,i<br>EDF 1a-e<br>EDF 2a,b<br>EDF 3e-g,o | 7 | 58 | 0, 0.5,<br>0.75, 1,<br>1.5, 2, 3,<br>4 | 0.5 | 2 |
| Varying delay<br>(5 delay types,<br>constant ITI duration) | Fig. 2e,f,l<br>EDF 1a-e<br>EDF 2a,b<br>EDF 3j-o | 7 | 72 | 0, 0.5,<br>0.75, 1,<br>1.5, 2, 3,<br>4 | 0.4, 0.5 | 1.6, 2 |
| Varying large amount | Fig. 2c,d,j<br>EDF 1f-j<br>EDF 2c,d | 7 | 36 | 0.5 | 0.2 | 0.4, 0.5, 0.8, 1,<br>1.2, 1.5, 1.6, 2 |
| Varying small amount | EDF 1k-o | 5 | 30 | 0.5 | 0.2, 0.5, 0.75,<br>1, 1.5, 2 | 2 |
| Varying delay and<br>large amount | Fig. 2g,h,k<br>Fig. 3<br>EDF 1p-t<br>EDF 2e-h<br>EDF 6<br>EDF 9e-h | 6 | 70 | 0, 0.5, 1,<br>1.25, 1.5,<br>2 | 0.4, 0.5 | 0.8, 1, 1.6, 2 |
| Varying delay<br>(3 delay types) | Fig. 4a-h<br>Fig. 5a-f<br>EDF 7<br>EDF 9a-d,<br>EDF 9i-n | 11 | 49 | 0, 0.5, 1,<br>1.25, 1.5 | 0.3, 0.4 | 1.5, 1.6 |
| MCx inhibition<br>with varying delay | Fig. 4i-k<br>EDF 8 | 10 | 10 | 0, 0.5,<br>1.25, 1.5 | 0.3, 0.4 | 1.6 |
| vPFC inhibition<br>with varying delay | Fig. 6<br>EDF 11a,e-h<br>EDF 12 | 10 | 10 | 0, 0.5, 1,<br>1.25, 1.5 | 0.3, 0.4 | 1.6 |
| vPFC inhibition<br>with varying amount | EDF 11b,i-l | 8 | 8 | 0.5 | 0.3, 0.4 | 0.8, 0.9, 1, 1.6 |
| ORB inhibition<br>with varying delay | EDF 11c,m-p | 6 | 6 | 0, 0.5, 1,<br>1.25, 1.5,<br>2 | 0.3, 0.4 | 1.6 |

